# Nitrogen uptake pattern of dry direct-seeding rice and its contribution to yield in northeastern Japan

**DOI:** 10.1101/2022.10.13.512177

**Authors:** Mari Namikawa, Takayuki Yabiku, Maya Matsunami, Toshinori Matsunami, Toshihiro Hasegawa

## Abstract

Dry direct-seeding rice (DDSR) cultivation is expected to reduce production costs compared with transplanted rice (TPR); however, its low nitrogen (N) use efficiency (NUE) has hindered cost reduction. Additionally, polymer-coated urea application in rice cultivation is reduced for plastic pollution regulation. The split application of urea can be an alternative, but it has not been used in northeastern Japan, hence needs to be investigated. We conducted DDSR and TPR field experiments for three years using two cultivars and three or two N regimes to determine factors limiting yield and NUE using a standard cultivar (‘Akitakomachi’) and a high-yielding cultivar (‘Yumiazusa’) grown under different N regimes. The yield, yield components, and N uptake of DDSR were analyzed, and examined the contribution of N uptake until panicle initiation and heading for spikelet number by multiple regression compared to that of TPR. Additionally, we investigated the detailed N uptake pattern on DDSR until PI using the two parameters, which were calculated by exponential regression of N uptake during the vegetative period. DDSR yield was lower than that of TPR by 11% and revealed that both fertilizer recovery rate and crop NUE (yield per unit N uptake) contributed to the lower yield. N uptake until the fifth leaf age significantly influenced the N uptake until panicle initiation. DDSR yield with normal urea in this study proportion was not significantly different compared to coated urea application, indicating the possibility to be an alternative N application method.

## Introduction

In Japan, the population involved in agriculture decreased substantially from 11.7 million in 1960 to 2.1 million in 2015, a reduction of 86% (MAFF, 2018), while increasing the cultivating land per farmer. This trend is projected to continue, and labor-saving and cost-effective technologies are expected to gain prominence. Dry direct-seeded rice (*Oryza sativa L.*) (DDSR) cultivation is thus becoming an important alternative for rice farmers in Japan since it can reduce labor costs compared to conventional transplanted rice (TPR) cultivation (Farooq et al., 2011).

Northeastern Japan is one of Japan’s main rice cultivation areas, producing approximately 29% of the national rice production (MAFF, 2021). This region has comparatively larger paddy fields than the rest of Japan, suitable for large-scale mechanized farming. Although both wet direct-seeded rice and DDSR technologies are available, DDSR has greater potential for saving labor than wet direct-seeded rice as larger agricultural machinery can be introduced for land preparation and planting practices (Kanmuri et al., 2017). DDSR technology, however, has some drawbacks that prevent it from being the more popular choice in northeastern Japan. First, DDSR yields are generally lower than TPR yields. MAFF (2021b) reported that direct-seeded rice, including dry-seeded and wet-seeded rice, yielded 7% less than TPR (the comparison of the average on 436 field data). Second, DDSR is associated with low nitrogen (N) use efficiency and requires greater fertilizer inputs (Farooq et al., 2011); thus, increasing production costs. Third, appropriate varieties for better performance in DDSR have not been studied in a cool climate.

Many of these problems are related to the climatic conditions in northeastern Japan, located in a cool temperate zone, where rice productivity has been severely limited by low temperatures (Horai et al., 2013). DDSR seedlings are exposed to low air temperatures from an earlier stage in contrast to the condition characteristic of TPR seedlings, which are covered or protected in nurseries such as greenhouses. One of the important advantages of ponding water is to protect rice plants from air cold temperatures because water is generally much warmer than air, particularly in the early stages of rice growth in cool temperate regions, which is known as the water blanket effect (Shimono et al., 2007). Shimono et al. (2007) demonstrated that in northern parts of Japan, simulated grain yields could be halved in the absence of surface water even without water stress because low temperatures at shoot bases hampered rice growth and yield. In DDSR, seedlings are grown under upland conditions without the water blanket for more than a month before permanent flooding. These situations possibly limit the yield potential of DDSR in a cool climate.

Temperatures are also highly influential on rice N uptake (Shoji et al., 1976; Takahashi et al., 1976), particularly during the vegetative growth stage (Shimono et al., 2012). The effects of N management on the growth and yield of DDSR have frequently been studied in warm areas such as northwest India, Southern U.S, and central China (Ali and Thind, 2015; Griggs et al., 2007; Liu et al., 2015; Mahajan et al., 2011; Qi et al., 2012; Xu et al., 2021) in the absence of cold temperature stresses. Still, evidence exists that DDSR needs a higher rate of N application than TPR for the same yield (Farooq et al., 2011; McDonald et al., 2006). However, little is known about rice N uptake during early-stage DDSR in cool climate zones. but N uptake of DDSR limited by low temperatures could be another reason for the lower N use efficiency (NUE).

The use of control-release urea is empirically recommended for DDSR in northern regions, and generally, a greater amount is recommended for DDSR than that for TPR (NARO, 2021). The fertilizer price of coated urea is approximately 3 times higher compared to normal urea per N g m^−2^, thus hindering the efforts toward reducing production costs. Additionally, polymer-coated urea uses microplastic capsules, which have recently been identified as a source of environmental pollution (Katsumi et al., 2020). Consequently, attempts to reduce the use of polymer-coated urea have already started in Japan and will likely expand in the future. Although the split application of urea might represent a low-cost alternative and has been used as a basic method globally, including for DDSR Europe and the USA, it needs to be investigated whether the split application of urea can be an effective option for DDSR in northeastern Japan.

High-yielding and good-quality cultivars are an important option for higher profitability; however, a comparison of cultivars for DDSR suitability in northeastern Japan has not been performed. In a cool climate, low temperatures limit growth duration, and DDSR without protected nursery periods takes longer to mature than TPR. Cultivars for DDSR, therefore, need to be early maturing, which is often not associated with high-yielding characteristics. Recently, however, an early-maturing, high-yielding, and good-eating quality cultivar, Yumiazusa, has been bred and released by NARO and is proven to be high-yielding for transplanting practice in cool temperate areas (Ohta et al., 2017; Yabiku et al, 2021).

Here, we examined how cultivar and N management combinations affect rice growth and yield under DDSR in a cool climate using a standard cultivar (‘Akitakomachi’, AK) and a high-yielding cultivar (‘Yumiazusa’, YA) grown under different N regimes. We focused on the following two points:

1. N uptake relevance on dry direct-seeded rice (DDSR) to yield and yield components compared to those of transplanted rice (TPR)
2. temperature-driven N uptake pattern analysis of DDSR for split application of normal urea (NU) compared to one-time polymer-coated urea (CU) application.

These results were compared with TPR experiments conducted side-by-side using the same cultivars (Yabiku et al., 2021).

## Material and methods

### Study sites

We conducted field experiments at the Experimental Farm of Tohoku Agricultural Research Center, NARO, in Morioka city, Iwate prefecture, Japan (39°45’N 141°08’E) in the 2018, 2019, and 2020 rice growing seasons. In each season, two fields were used for the experiments: one for DDSR and the other for TPR. The soils of all experimental fields were classified as Andisols, and some of their chemical properties are shown in Table S1.

Climatic data were obtained from the meteorological observation at the NARO Tohoku Agricultural Research Center located 2 km from the experimental fields. Table S1 presents mean air temperatures and mean precipitations from April to October of the experimental years (2018–2020).

### Experimental design and field management

#### Plant materials and N treatments

We used two temperate *japonica* cultivars: ‘Akitakomachi’ (AK) and ‘Yumiazusa’ (YA). AK is an old but still widely planted cultivar in northeast Japan, whereas YA is a high-yielding cultivar with high eating quality, released in 2017 (Ohta et al. 2017). For DDSR, we had three N regimes: two with N fertilizers and one without (0N). In the two fertilized plots, polymer-coated urea (CU) and normal urea (NU) were used. For the CU treatment, we applied 12 g N m^−2^ of CU fertilizer as basal, containing three types of slow-release fertilizer differing in the rate and pattern of N release (LP30:30%, LPS30: 20%, LPS60: 50%, Chokuha senyou 211, Kumiai Hiryou Co. Ltd., Iwate, Japan). For the NU treatment, a total of 15g N m^−2^ of urea was split-applied four times at 40, 66, 82, and 94 days after sowing (DAS) in 2018, 45, 66, 80, 94 DAS in 2019, and 52, 67, 80, 95 DAS in 2020, at a dose of 6 g N m^−2^, 3 g N m^−2^, 3 g N m^−2^, and 3 g N m^−2^, respectively. In the 0N treatment, no N fertilizer was applied. Equal amounts of P and K were applied to all plots at the rate of 3.4g P m^−2^ and 3.1g K m^−2^ as basal, respectively, on 23 April (2018), 17 April (2019), and 15 April (2020).

All experiments in DDSR were laid out in a randomized complete block design, where a factorial combination of two cultivars (AK and YA) and three N regimes (0N, CU, and NU) was randomly assigned to each of the three blocks. The size of each plot was 192 m^2^ (40 m × 4.8 m, CU treatment) or 48m^2^ (20 m × 2.4 m, 0N, and NU treatment) in 2018, and 81 m^2^ (15 m × 5.4 m) or 124 m^2^ (23 m × 4.8 m) in 2019 and 2020.

In TPR, the same cultivars were grown under three different N doses: 0, 8, and 18 g m^−2^, denoted as 0N, 8N, and 18N, respectively in 2018-2020 (details in 2018 are presented in Yabiku et al. (2021)). 8N was considered a standard N level for TPR, comparable to the CU treatment for DDSR. 18N was used to only analyze the relationship between crop N uptake at maturity and yield components. The N, P, and K were applied as basal by compound fertilizer (Iseki court M002, Iseki Tohoku, Miyagi, Japan) in 8N and 18N, and P and K were applied to all plots at the rate of 3.5 g P m^−2^ and 4.0g K m^−2^. 18N plots were top-dressed 2 g N m^−2^ of ammonium sulfate and 4 g N m^−2^ of coated urea (LP40, JCAM AGRI. Co. Ltd., Tokyo, Japan) at the tillering stage, 2 g N m^−2^ of ammonium sulfate at the panicle initiation stage, 4 g N m^−2^ coated urea (LP30, JCAM AGRI. Co. Ltd., Tokyo, Japan) and 3.3 g K m^−2^ potassium chloride at spikelet differentiation stage. The N treatments and cultivars were laid out in a split-block design for three years with three replicated each year.

#### Cultivation details

In DDSR, the dry soil was plowed, power-harrowed, and laser-leveled in April. Basal fertilizer was applied approximately three days before sowing. After basal fertilizer application, the soil was tilled with a vertical harrow tiller and compacted using a Cambridge roller. Before sowing, seeds were coated with a thiram-based repellent (bactericide) (Kihigen R-2 flowable, Yonezawa Chemical Co., Ltd., Kyoto, Japan). We sowed the dried seeds to a depth of approximately 2 cm using the high-speed seeder (Clean seeder; NTP-6AFP and NTP-8AFP, Agritechno Search Co. Ltd, Hyogo, Japan) at a row spacing of 0.3 m on 27 April (2018), 22 April (2019), and 17 April (2020). Immediately after seeding, the lands were suppressed by Cambridge roller for good seedling establishment and impermeability (Kanmuri et al., 2017). The sowing rate was 5.1 g m^−2^ (2018), 5.7 g m^−2^ (2019), and 5.7 g m^−2^ (2020). We applied flush irrigation about once in three days to prevent the lands from drying until seedling establishment. From early June to approximately three weeks before harvest (late September), the field was kept flooded with approximately 3–10 cm ponding water.

Before permanent flooding, we applied glyphosate-potassium at a rate of 4.8 mL m^−2^ before emergence (mid-May) and cyhalofop-butyl (0.3 mL m^−2^), and bentazone (2 ml m^−2^) after emergence (early June), using a boom sprayer. In late June (after flooding), we applied a compound herbicide containing pyriftalid, mesotrione, and metazosulfuron at a rate of 90 mg m^−2^, 240 mg m^−2^, and 80 mg m^−2^ using a backpack spreader.

For TPR, the dry soil was plowed in mid-April, and basal fertilizers were broadcast on the soil surface and incorporated into the soil with the rotary tractor during early May. During mid-May, the field was submerged and puddled with a power harrow system. Seedlings were raised in a greenhouse and at about fifth plant age in leaf number (leaf age), seedlings were mechanically transplanted on May 17, 21, and 19 in 2018, 2019, and 2020, respectively with a spacing of 30 × 19 cm. The field was kept flooded from May until the end of August, after which the surface water was withdrawn for harvest in late September.

### Plant measurements, sampling, and tissue N determination

Leaf age was measured on the three plants in each replicate at one-week intervals from emergence to flag leaf development. At the same time, tillers were counted on 0.3 m^2^ (one 1 m row). From approximately 45 DAS until harvest, we sampled aboveground plant organs weekly or bi-weekly, washed, oven-dried them at 80 °C for at least 72 hours, and weighed them to determine the aboveground biomass. The sampled area at each harvest was 0.3 m^2^ (one 1 m row). Part of the dried samples was ground, and tissue N concentrations were measured using an elemental analyzer (Variomax, Elementar, Germany). Aboveground crop N uptake (*N_up_*) was calculated by multiplying the biomass and tissue N concentrations.

At harvest, plants were collected from an area of 2.4 m^2^ (1.2m (4rows) × 2m) in each plot for yield determination. We air-dried the samples, weighed the total mass, threshed and dehulled the grains, and screened them through a 1.85 mm-thick sieve. Straw and grain moisture contents were also measured. Brown rice yield was expressed on the measured sample for yield determination. The water content of brown rice samples was corrected to 15%.

To analyze N use efficiency, Agronomic NUE (ANUE) in the fertilized plots was calculated as:

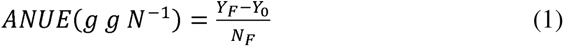

where *Y_F_* and *Y_0_* are brown rice yields from the fertilized and 0N plots, respectively, and *N_F_* is the amount of N fertilizer applied. ANUE can be considered a product of fertilizer N recovery and the efficiency of converting crop N into grain yield.

The apparent N recovery (ANR) in the plots was defined as:

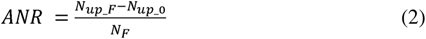

where *N_up_F_* and *N_up_0_* are *N_up_* in the fertilized plot and 0N plot, respectively.

We also derived crop NUE (CNUE), which was defined as yield per unit of N uptake at maturity and partial CNUE fertilization (CNUEF) as:

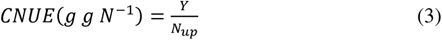

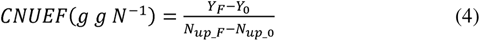

ANUE can be derived by multiplying ANR and CNUEF: ANUE = ANR×CNUEF (Eqs 1, 2, and 4).

### Analysis of N uptake during the vegetative growth stage

N uptake is closely related to temperature conditions (Shimono et al., 2012) and is known to be approximated by the function of cumulative air temperature. In transplanted rice, Takahashi et al. (1976) showed that the accumulated effective thermal index (AETI) could reproduce well the N uptake pattern under different conditions (cropping seasons or years), which is defined as

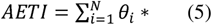

where θ_i_ * is the daily effective thermal index, which is the product of the hourly air temperature and *effective air temperature coefficient* (Table S2), averaged for the i-th day (Hanyu and Uchijima 1962), and N is the number of days from start to end of the calculation. In this study, we set i=1 at the fifth leaf age, and the exact date when the leaf age becomes 5.0 was derived from the linear interpolation of the weekly observations. The hourly air temperatures (*T_j_*) were approximated using the following cosine curve (Sun et al., 2018):

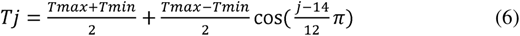

where *T_max_* and *T_min_* are the daily maximum and minimum air temperature, respectively, *j* is the time of day, and *T_j_* is the hourly air temperature at time *j*.

Previous studies showed that N uptake has two distinctive phases (Shoji et al., 1976; Takahashi et al., 1976). In phase 1, *N_up_* exponentially increases, limited by the crop growth rate, and can be expressed as an exponential function of AETI:

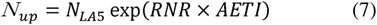

where *N_LA5_* is *N_up_* when leaf age is 5.0 (LA5), and *RNR* is the relative *N_up_* rate. In phase 2, *N_up_* increases linearly, limited by soil N supply (not analyzed in this paper).

Accordingly, *N_up_* for the vegetative growth period can be characterized by two parameters: *N_LA5_*, and *RNR*. To derive these parameters, we first determined the duration of the exponential phase from LA5 to the panicle initiation date in each cultivar and year. We then exponentially regressed *N_up_* on AETI to obtain *N_LA5_* and *RNR*. All parameters were calculated in each replicate in 2018 and 2019 and provided for the statistical analysis (next section).

### Statistical analysis

Statistical analyses were performed using JMP12.0.1 (SAS Institute, 2015). *P* values < 0.05 were considered significant. We conducted an analysis of variance (ANOVA) for DDSR yield components and N uptake parameters using a split-plot analysis, where the year was treated as the main plot, and cultivar and N treatment as the subplots. The model included three fixed effect variables (Year (Y), cultivar (C), and N treatments (N)) and one random effect variable (block) with the interactions (Y*C, C*N, Y*N, Y*C*N). Tukey’s HSD test was performed on some effects of ANOVA, with a significant level of *P*<0.05.

Yield and yield components are related to plant N uptake. We computed the analysis of covariance (ANCOVA) for brown rice yield and each yield component as objective variables, N uptake at maturity stage (*N_MT_*) as explanatory variables, and planting method (DDSR or TPR, dummy variables) as covariance. 81 plots on DDSR and 54 plots on TPR of each experiment were used on ANCOVA.

Although N uptake before the heading stage is a critical determinant for spikelet density (Hasegawa et al., 1994; Sasaki, 2007), the N regimes may alter the relative contribution of N uptake at different stages. We, therefore, applied multiple regression analysis for spikelet density as a function of N uptake at panicle initiation (*N_PI_*) and N uptake from then to heading (*N_PI-HD_*). Multiple regression of Model 1 was performed with spikelet density as an objective variable, *N_PI_*, N uptake at the heading (*N_HD_*), or *N_PI-HD_*, and additional categorical variables (*Planting method* (DDSR or TPR), *Cultivar* (‘Akitakomachi’ or ‘Yumiazusa’), and *Year* (2018, 2019, or 2020)). Model 2 was similarly conducted, however, not considered categorical variables (*Planting method*, *Cultivar*, and *Year*).

*N_LA5_* and *RNR* are indicators of N uptake during the vegetative growth period. We investigated the contribution of initial N uptake and increase rate during the vegetative growth period for N_PI_: multiple regression analysis was applied for N_PI_ using *N_LA5_* and *RNR* including or excluding categorical variable of *Year* (2018 and 2019). Model 3 included and Model 4 excluded *Year*.

The standard least-squares method was used as a fitting methodology. The adjusted coefficient of determination (Adjusted R^2^), corrected Akaike’s Information Criterion (AICc), and Bayesian Information Criterion (BIC) were obtained to compare the two models. Adjusted R^2^ is the R square adjusted by following equation to be able to compare the two different regression models with a different number of explanatory variables.

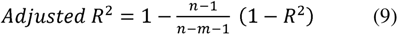

Where *n* is the sample size and *m* is the number of explanatory variables. The standardized partial regression coefficient (β) was also computed to determine which parameter has the higher impact on N_PI_ in each regression parameter of the selected model. Variance inflation factor (VIF) was used to consider multicollinearity among the explanatory variables.

## Results

### Meteorological data

In April-October 2018, 2019, and 2020, mean air temperatures were 0.42°C, 0.92°C, and 0.62°C higher than the 30-year mean (Table S1). In 2019, the mean air temperature in May was 2.1 °C higher than the 30-year mean, whereas these differences from the 30-year mean in 2018 and 2020 were within 1.0 °C.

### Crop development of DDSR

Emergence occurred at 18–21 DAS, and plants reached the second leaf stage (LA2) at 21–27 DAS in all DDSR experiments (Table S3). Dates of panicle initiation (PI) and heading (HD) in DDSR were later than that of TPR and similar across different environments and cultivars, ranging from 91–96 DAS and 114–120 DAS, respectively. The period from heading to maturity tended to be longer for YA than for AK. We reported the results of tiller number and biomass growth in the supplemental materials (Figures S3 and S4).

Leaf age growth was impacted by the N regimes from the early stages (approximately 50 DAS; Figure S1). Days from sowing to LA5 were 2-10 days earlier in CU than in 0N and NU (Table 1). The difference in leaf age between 0N and CU persisted until the final leaf development, (0.4–1.7 fewer total leaves on the main culm in 0N than in CU (Figure S2)). This difference was approximately 65% accounted for by the difference at the seedling stage (Figure S2, the slope of regression on the relationship between the leaf number differences in the CU treatment condition at the seedling stage and the final leaf number). Leaf age increase in the NU treatment was initially slow but recovered from 50 DAS (Figure S2), narrowing the gap with CU to approximately 0.25 averaged at the end of the experiment (Table 1).

**Table 1.** The differences on leaf ages at each stage and days after sowing among N regimes in DDSR.

| Year | Cultivars | N regimes | LA2 <sup>1</sup><br>DAS | LA5 <sup>2</sup><br>DAS | The leaf ages<br>at the seedling<br>stage <sup>3</sup> | The final<br>leaf number |
| --- | --- | --- | --- | --- | --- | --- |
| 2018 | AK | 0N | 27 | 54 | 4.6 | 13.3 |
|  |  | CU | 27 | 47 | 4.8 | 14.2 |
|  |  | NU | 27 | 49 | 4.8 | 14.1 |
|  | YA | 0N | 27 | 49 | 4.8 | 13.7 |
|  |  | CU | 27 | 45 | 5.1 | 14.5 |
|  |  | NU | 27 | 48 | 4.7 | 13.9 |
| 2019 | AK | 0N | 24 | 57 | 4.2 | 11.4 |
|  |  | CU | 24 | 47 | 5.4 | 13.1 |
|  |  | NU | 24 | 51 | 4.7 | 13.1 |
|  | YA | 0N | 24 | 55 | 4.3 | 11.6 |
|  |  | CU | 24 | 48 | 5.2 | 12.7 |
|  |  | NU | 24 | 53 | 4.4 | 12.4 |

DAS, days after sowing.
AK and YA represent the cultivar's names, 'Akitakomachi' and 'Yumiazusa'.
0N, CU, NU, and 8N represent the N treatments, no N treatment, one-time coated urea treatment (12 g N m<sup>-2</sup>), split application of normal urea (15 g N m<sup>-2</sup>, 6, 3, 3, 3), and conventional compound fertilizer treatment (8 g N m<sup>-2</sup>), respectively.
<sup>1</sup> LA2, 2<sup>nd</sup> leaf age (perspective survey).
<sup>2</sup> LA5, 5<sup>th</sup> leaf age.
<sup>3</sup> The seedling stage was at 45 (2018), and 49 (2019) days after sowing, respectively.

### Yield and yield components

DDSR brown rice yield under CU conditions was 452 g m^−2^ averaged for the three years and two cultivars, 89% of the average yield of transplanted rice under 8N conditions of the same varieties (Table 2, Table S4, and Table S5). DDSR brown rice yield differed significantly among years, with the 2019 crop showing a considerably lower yield than the 2018 and 2020 crops (*P* = 0.0174, Table S5). The yields of CU and NU were not significantly different on Tukey’s HSD test; however, the NU/CU ratio tended to be lower than 1.0 (Table 2). Of the two cultivars on DDSR, YA had an 8% greater yield than AK, averaged over treatments and years (*P* = 0.0083), although the cultivar effect was not consistent among years (*P* = 0.0062 for Y * C interaction, Table 1 and Table S5). The yield advantage of YA over AK was apparent in relatively high-yielding years (16% in 2018 and 12% in 2020, Table S5), while AK had a higher yield in 2019 averaged over the three N regimes.

**Table 2.**
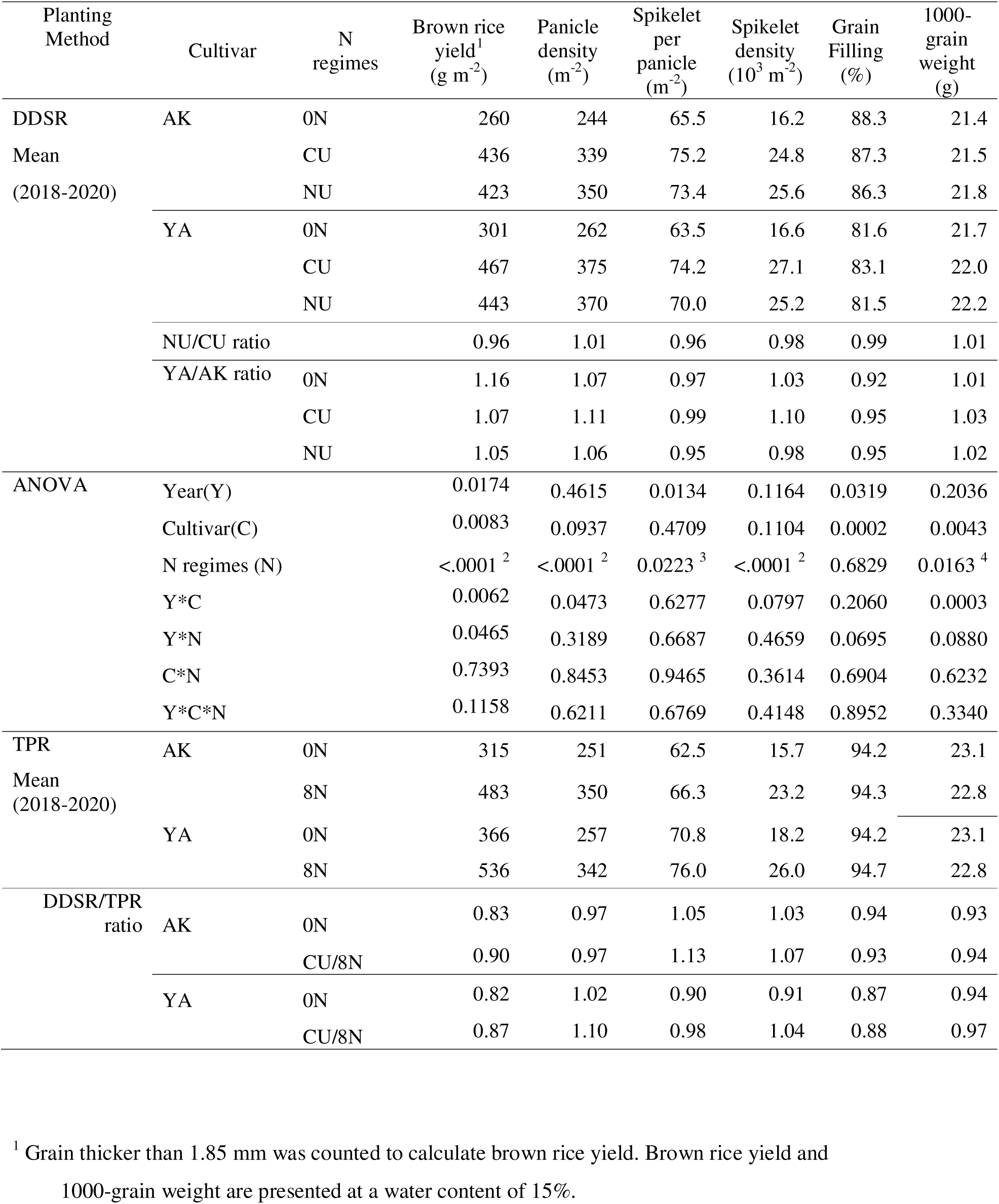
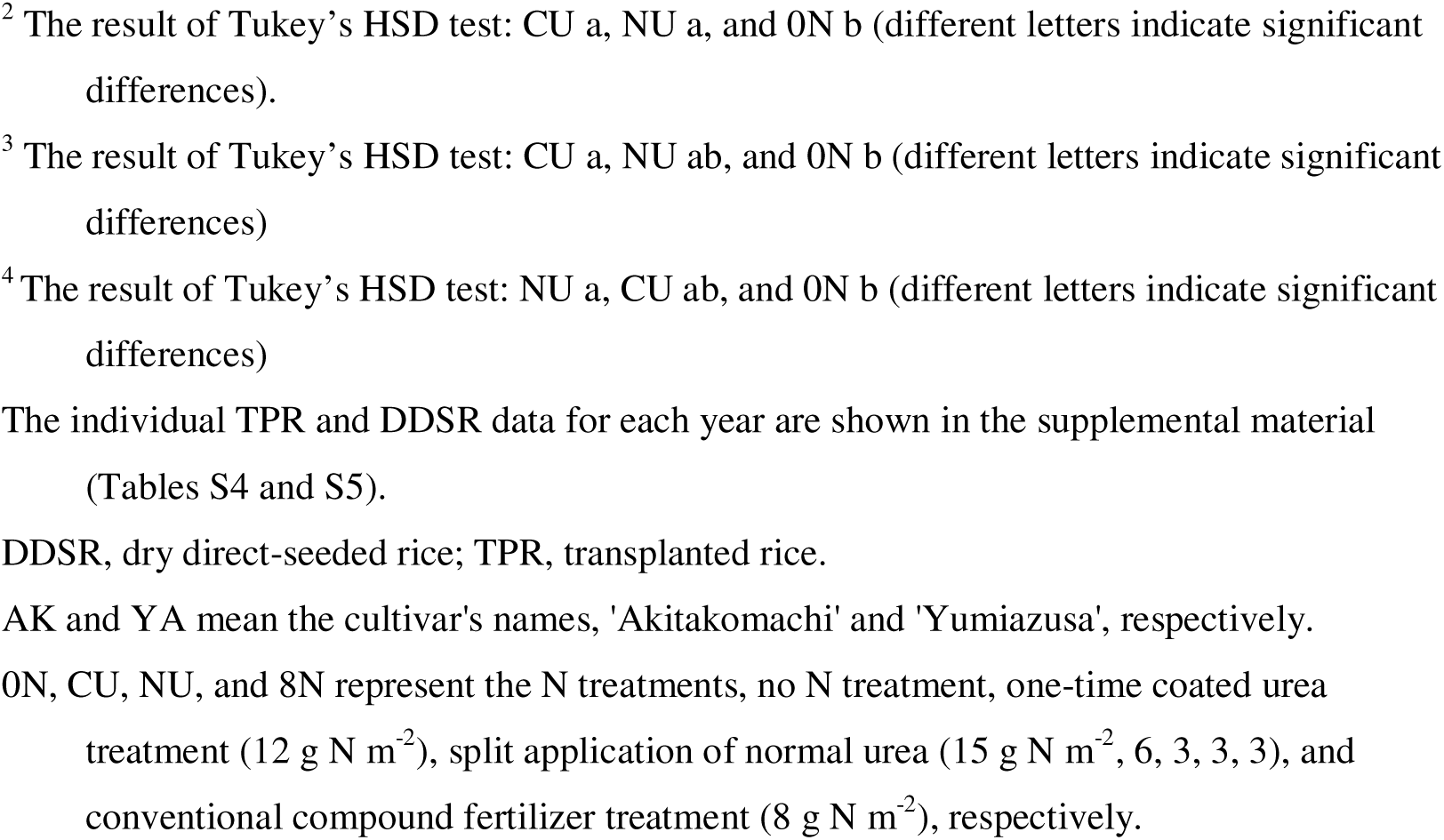
Brown rice yield and yield components under different nitrogen regimes and cultivars.

| Planting Method | Cultivar | N regimes | Brown rice yield<sup>1</sup><br>(g m<sup>-2</sup>) | Panicle density<br>(m<sup>-2</sup>) | Spikelet per panicle<br>(m<sup>-2</sup>) | Spikelet density<br>(10<sup>3</sup> m<sup>-2</sup>) | Grain Filling (%) | 1000-grain weight (g) |
| --- | --- | --- | --- | --- | --- | --- | --- | --- |
| DDSR<br>Mean<br>(2018-2020) | AK | 0N | 260 | 244 | 65.5 | 16.2 | 88.3 | 21.4 |
|  |  | CU | 436 | 339 | 75.2 | 24.8 | 87.3 | 21.5 |
|  |  | NU | 423 | 350 | 73.4 | 25.6 | 86.3 | 21.8 |
|  | YA | 0N | 301 | 262 | 63.5 | 16.6 | 81.6 | 21.7 |
|  |  | CU | 467 | 375 | 74.2 | 27.1 | 83.1 | 22.0 |
|  |  | NU | 443 | 370 | 70.0 | 25.2 | 81.5 | 22.2 |
|  | NU/CU ratio |  | 0.96 | 1.01 | 0.96 | 0.98 | 0.99 | 1.01 |
|  | YA/AK ratio | 0N | 1.16 | 1.07 | 0.97 | 1.03 | 0.92 | 1.01 |
|  |  | CU | 1.07 | 1.11 | 0.99 | 1.10 | 0.95 | 1.03 |
|  |  | NU | 1.05 | 1.06 | 0.95 | 0.98 | 0.95 | 1.02 |
| ANOVA | Year(Y) |  | 0.0174 | 0.4615 | 0.0134 | 0.1164 | 0.0319 | 0.2036 |
|  | Cultivar(C) |  | 0.0083 | 0.0937 | 0.4709 | 0.1104 | 0.0002 | 0.0043 |
|  | N regimes (N) |  | <.0001<sup>2</sup> | <.0001<sup>2</sup> | 0.0223<sup>3</sup> | <.0001<sup>2</sup> | 0.6829 | 0.0163<sup>4</sup> |
|  | Y*C |  | 0.0062 | 0.0473 | 0.6277 | 0.0797 | 0.2060 | 0.0003 |
|  | Y*N |  | 0.0465 | 0.3189 | 0.6687 | 0.4659 | 0.0695 | 0.0880 |
|  | C*N |  | 0.7393 | 0.8453 | 0.9465 | 0.3614 | 0.6904 | 0.6232 |
|  | Y*C*N |  | 0.1158 | 0.6211 | 0.6769 | 0.4148 | 0.8952 | 0.3340 |
| TPR<br>Mean<br>(2018-2020) | AK | 0N | 315 | 251 | 62.5 | 15.7 | 94.2 | 23.1 |
|  |  | 8N | 483 | 350 | 66.3 | 23.2 | 94.3 | 22.8 |
|  | YA | 0N | 366 | 257 | 70.8 | 18.2 | 94.2 | 23.1 |
|  |  | 8N | 536 | 342 | 76.0 | 26.0 | 94.7 | 22.8 |
| DDSR/TPR<br>ratio | AK | 0N | 0.83 | 0.97 | 1.05 | 1.03 | 0.94 | 0.93 |
|  |  | CU/8N | 0.90 | 0.97 | 1.13 | 1.07 | 0.93 | 0.94 |
|  | YA | 0N | 0.82 | 1.02 | 0.90 | 0.91 | 0.87 | 0.94 |
|  |  | CU/8N | 0.87 | 1.10 | 0.98 | 1.04 | 0.88 | 0.97 |

<sup>1</sup> Grain thicker than 1.85 mm was counted to calculate brown rice yield. Brown rice yield and 1000-grain weight are presented at a water content of 15%.
<sup>2</sup> The result of Tukey's HSD test: CU a, NU a, and 0N b (different letters indicate significant differences).
<sup>3</sup> The result of Tukey's HSD test: CU a, NU ab, and 0N b (different letters indicate significant differences)
<sup>4</sup> The result of Tukey's HSD test: NU a, CU ab, and 0N b (different letters indicate significant differences)
The individual TPR and DDSR data for each year are shown in the supplemental material (Tables S4 and S5).
DDSR, dry direct-seeded rice; TPR, transplanted rice.
AK and YA mean the cultivar's names, 'Akitakomachi' and 'Yumiazusa', respectively.
0N, CU, NU, and 8N represent the N treatments, no N treatment, one-time coated urea treatment ( $12 \text{ g N m}^{-2}$ ), split application of normal urea ( $15 \text{ g N m}^{-2}$ , 6, 3, 3, 3), and conventional compound fertilizer treatment ( $8 \text{ g N m}^{-2}$ ), respectively.

Panicle density and spikelet density were comparable between CU and NU (Table 2), although leaf age (Figure S1) and tiller number (Figure S3) progress differed between the two fertilized plots. The NU/CU ratio of panicle and spikelet density were smaller in 2018 than in the other two years (Table S5). Conversely, the NU/CU ratio for 91% of ripened spikelets and 1000-grain weight was greater in 2018, thus compensating for the limited sink size under NU conditions. The N regime significantly affected 1000-grain weight (*P* = 0.0163, Table 2), with NU having the heaviest mass, followed by CU and 0N.

The yield advantage of YA over AK was associated mainly with the greater panicle density (8%) averaged over years and N regimes. YA had slightly fewer spikelets per panicle and Grain filling than AK, but these differences did not offset the yield advantage of YA. The cultivar differences in these yield components were generally consistent across different N regimes and years (Table S5).

### N use efficiency and crop N uptake at key growth stages

N_MT_ of DDSR was 7.9 g N m^−2^, 11.2 g N m^−2^, and 11.8 g N m^−2^ in 0N, CU, and NU averaged over three years and two cultivars (*P* < 0.0001, Table 3). N_MT_ of DDSR were similar to or even greater than crop N uptake observed in TPR plants undergoing comparable N treatments (Figure 2). Furthermore, N_PI_ and N_MT_ in DDSR were no significant differences between cultivars, although N_HD_ had a significant Y*C effect (Table 3). N_HD_ in DDSR tended to be lower than that of TPR, whereas N_PI_ was lower at both 0N and fertilized plots than that of TPR (Figure 2). This resulted in a greater N uptake between PI and HD in the DDSR than in TPR plants. In the two DDSR fertilized plots, CU enhanced N uptake by PI more than NU, while NU overtook CU before HD (Figure 2).

**Table 3.**
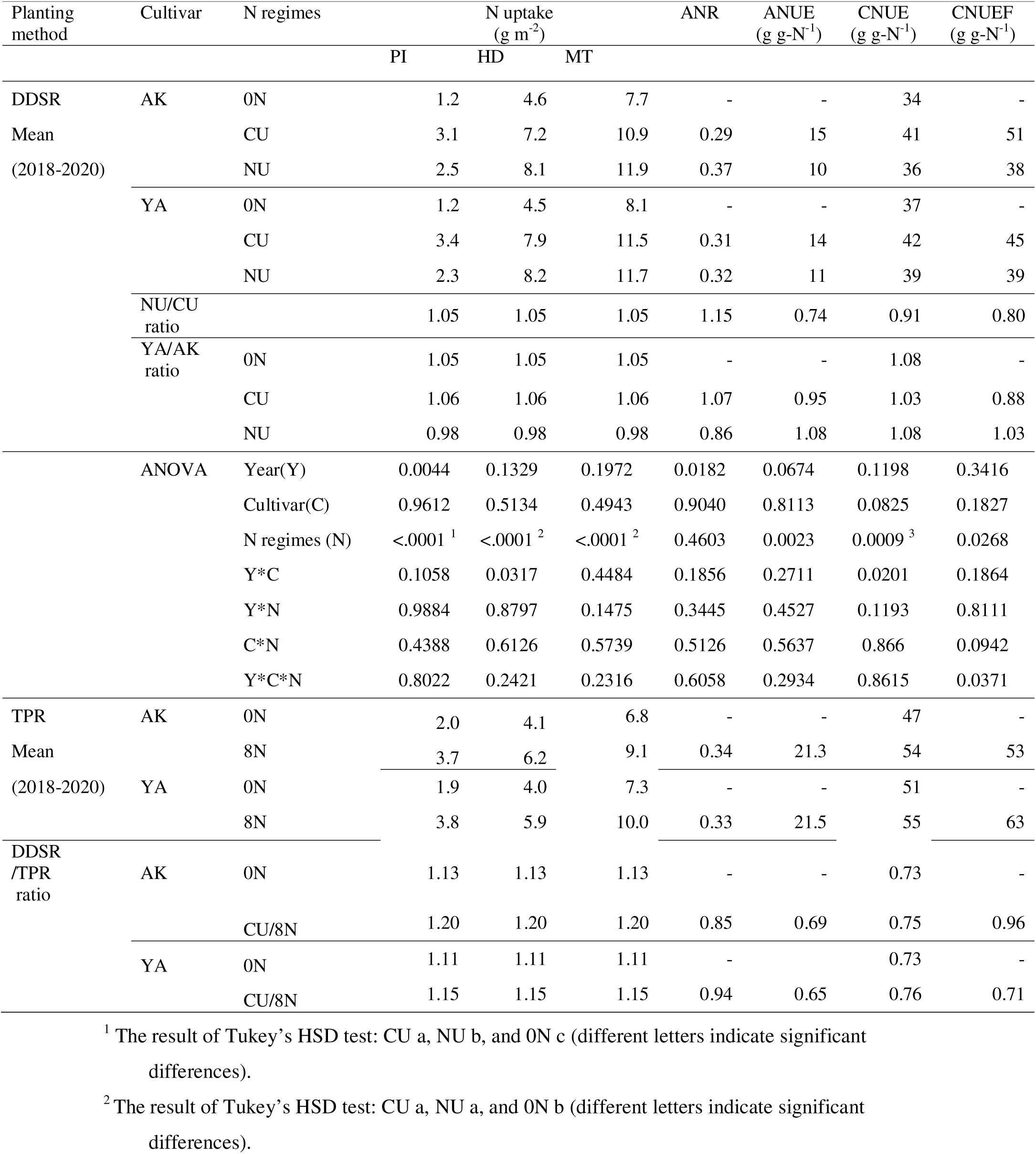
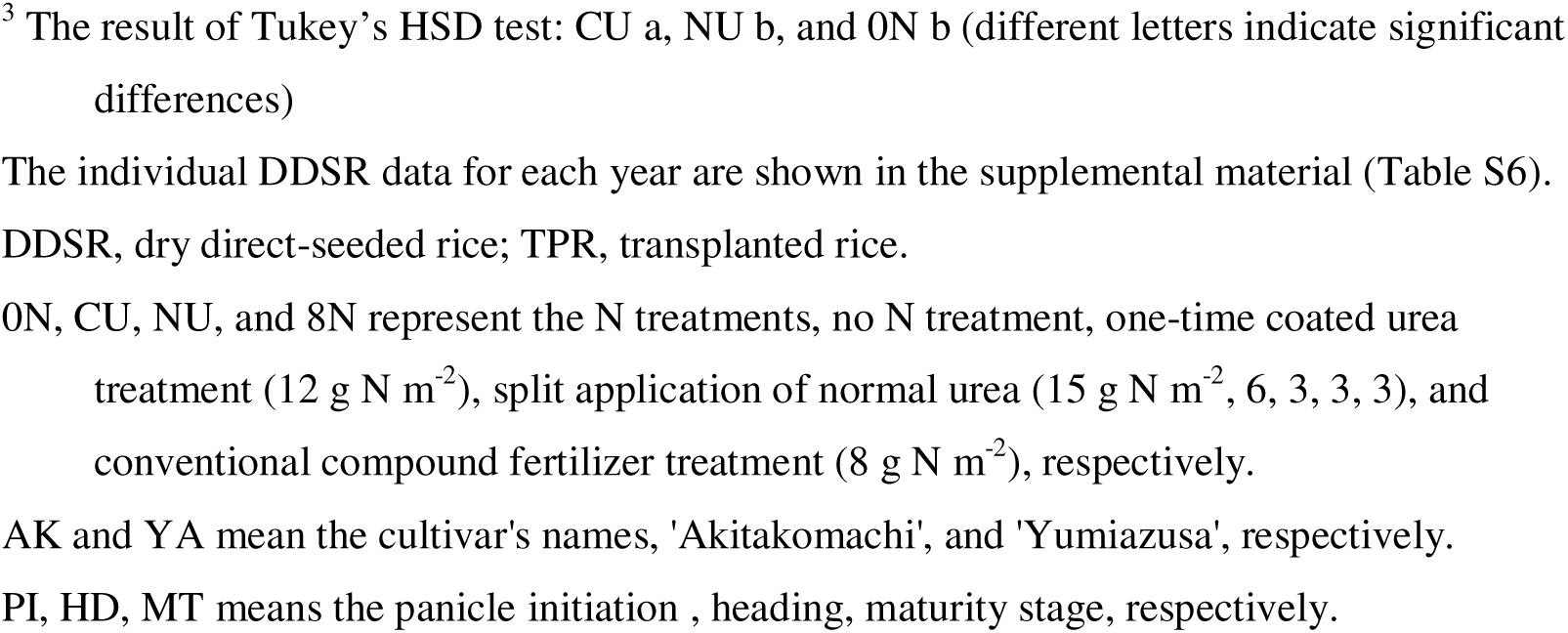
Rice plant nitrogen uptake and apparent nitrogen recovery rate of dry-direct seeded rice under different nitrogen regimes and cultivars.

The ANR was significantly different between years (*P* = 0.0182; Table 3), with lower values in 2018 than in the other two years (Table S6). The ANR was similar between CU and NU (*P* = 0.4603), averaging 0.30 in CU and 0.35 in NU for the three years, 90% in CU and 103% in NU relative to the 8N conditions of TPR (Table 3, Table S6). Conversely, ANUE was significantly lower in the NU than in the CU (*P* = 0.0023; 24% decrease) in both cultivars. The DDSR/TPR ratio for ANUE was 0.68 when averaged for CU (Table 3). CNUE was significantly different between the N treatment (*P* = 0.0012; Table 3, Table S6), and that of the CU treatment was significantly higher than that of the 0N and NU treatments (Tukey’s HSD test, *P*<0.05). CNUEF, the criteria of NUE that excluded soil N supply, showed an interaction with the year, cultivar, and N treatment variables. However, the CNUEF of the CU was consistently higher than that of the NU (Table 3).

### Relationship between crop N uptake at maturity and yield components

We examined the relationships between N_MT_ and yield components and compared them between DDSR and TPR to determine whether crop N utilization for the yield formation process differs by planting methods (Figure 1). The brown rice yield increased almost linearly with N_MT_, but the intercept was significantly greater for TPR than DDSR, confirming that DDSR had a lower CNUE than TPR (Figure 1a). Similarly, spikelet density had a strong positive relationship with N_MT_ in both TPR and DDSR, while spikelet density tended to be lower in DDSR than in TPR for the same N_MT_ (Figure 1b). Grain filling (%) in DDSR was not correlated with N_MT_ and was consistently lower than that in TPR (Figure 1c). The 1000-grain weight in DDSR was not modeled by N_MT_, planting method, and their interaction (R^2^ = 0.2982; Figure 1d).

**Figure 1.**
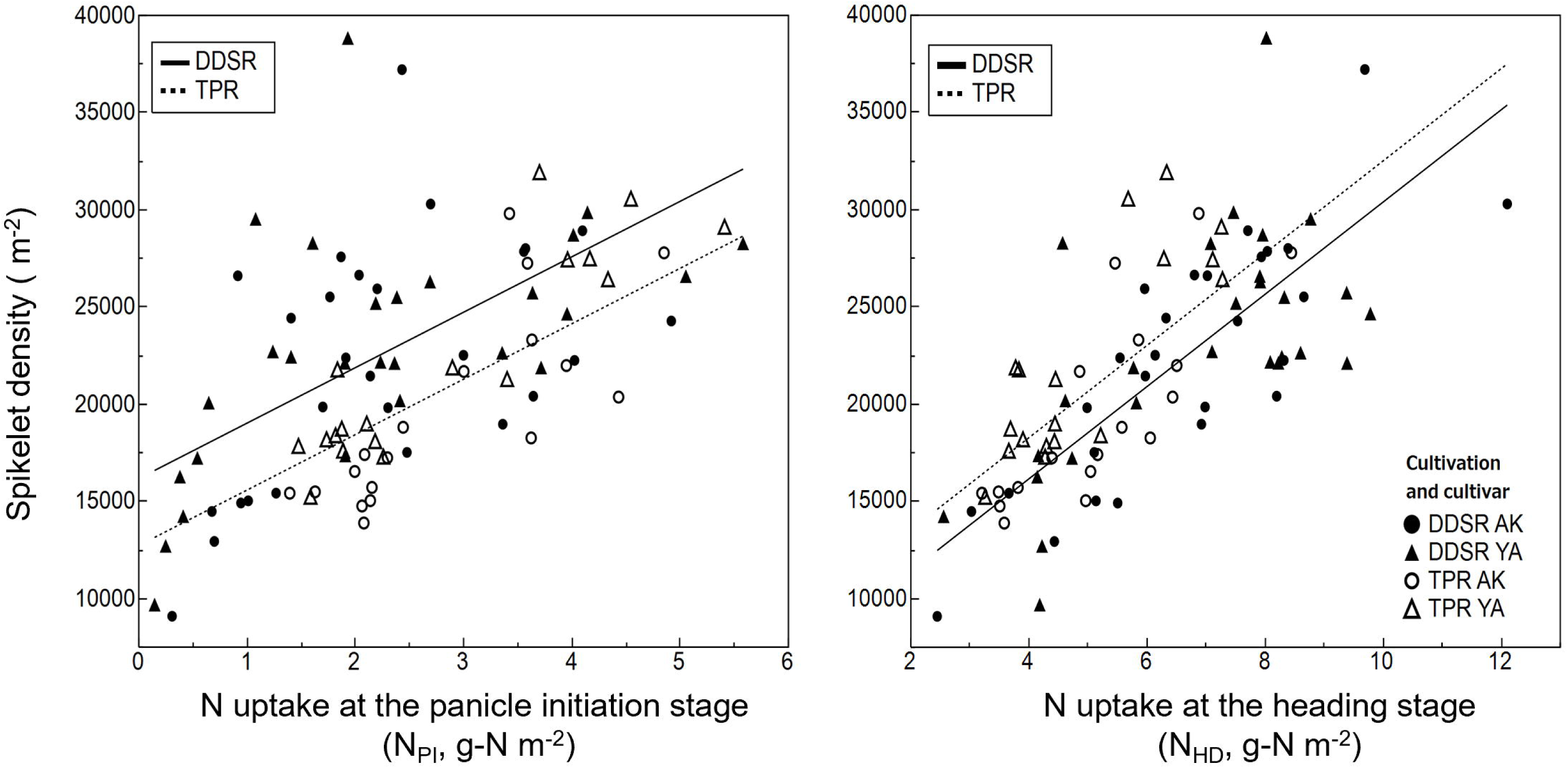
The relationship between N uptake at maturity (N_MT_) and yield components. From the analysis of covariance, the regression intercept for brown rice yield was different between TPR and DDSR (*P* < 0.0001; (a)). The spikelet density relationship with N_MT_ was different as a function of the planting method (*P* < 0.0001; (b)). The regression intercept was different for grain-filling (%) between TPR and DDSR (*P* < 0.0001; (c)). The 1000-grain weight was not influenced by N_MT_, planting method, and that interaction (R^2^ = 0.2982, (d)).

**Figure 2.**
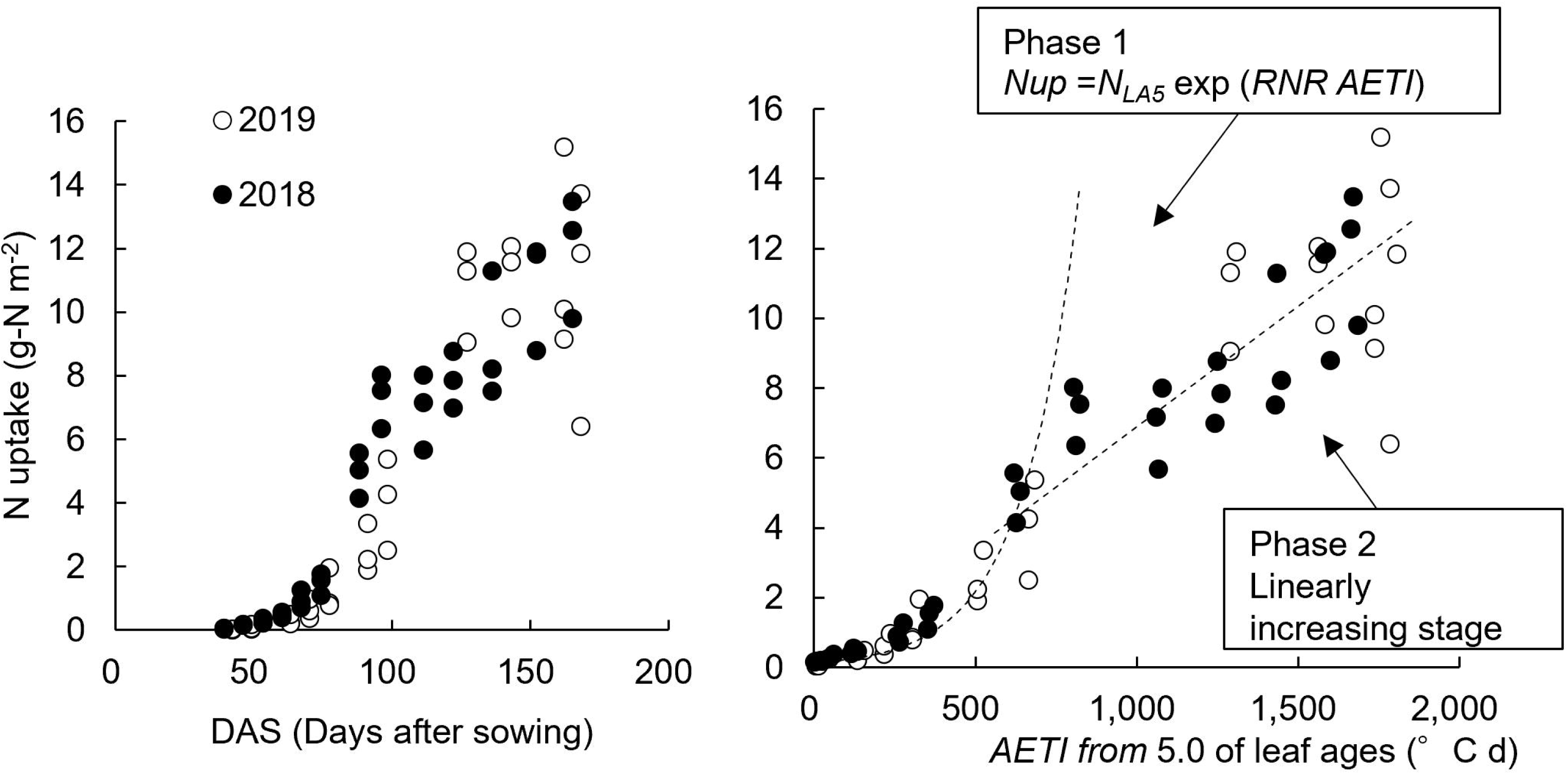
The N uptake for each N treatment, planting method, and cultivar. 0N, CU, NU, and 8N represent the N treatments, no N treatment, one-time coated urea treatment (12 g N m^−2^), split application of normal urea (15 g N m^−2^, 6, 3, 3, 3), and conventional compound fertilizer treatment (8 g N m^−2^), respectively. DDSR and TPR represent the dry direct seeded rice and transplanted rice, respectively.

### Analysis of the relationships between N uptake and spikelet density

Spikelet density was also significantly correlated with N_PI_ and N_HD_, and it was observed that the regression lines were different between TPR and DDSR at both stages (Figure 3, significantly different intercepts; *P* = 0.002). However, there was a clear difference between the two stages, where the regression line for DDSR was higher than that for TPR at PI, and it was reversed at HD (Figures 3a, b).

**Figure 3.**
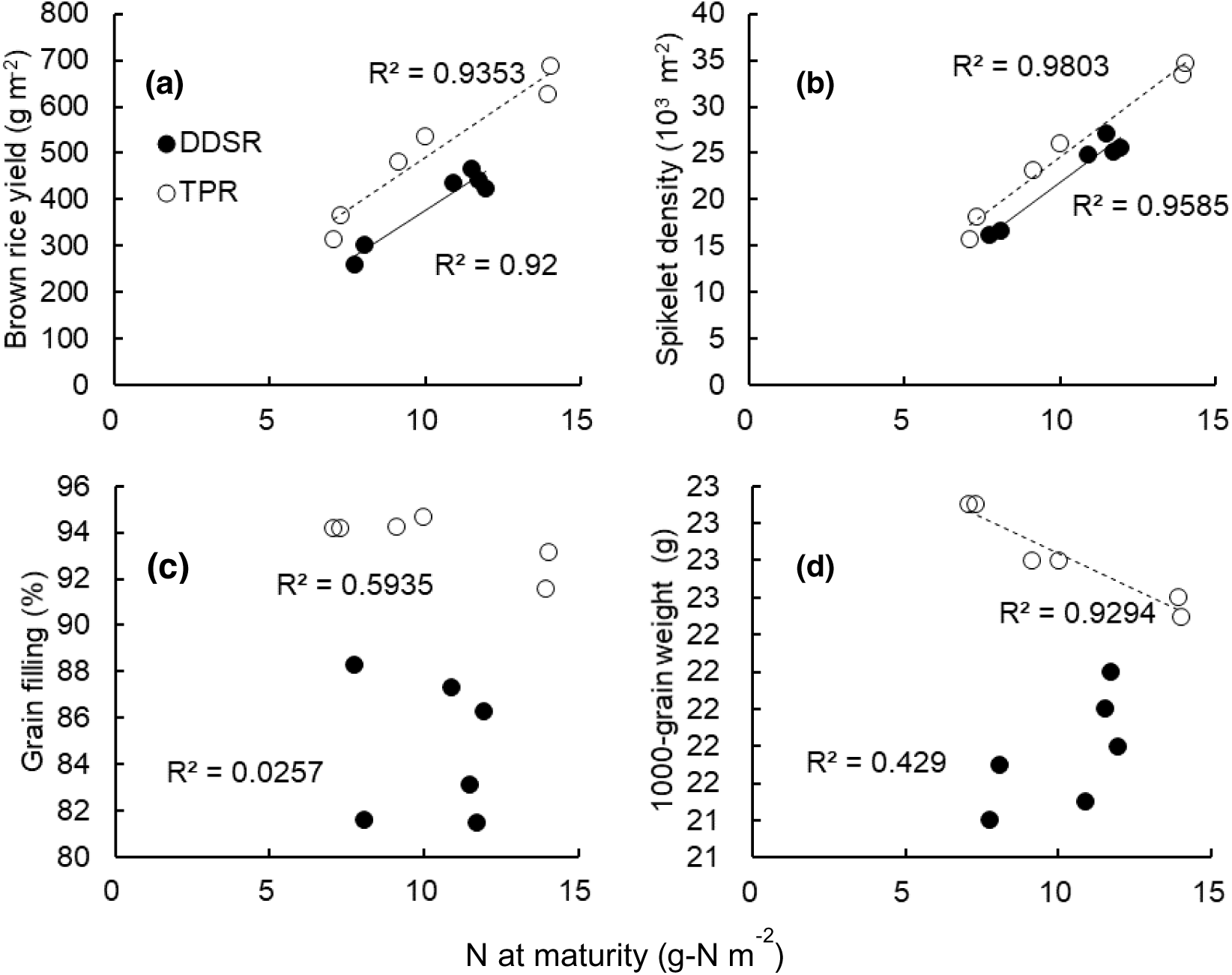
The relationship between spikelet density and N uptake at *N_PI_* or *N_HD_* under different planting methods, DDSR and TPR. The difference relationship in *N_HD_* was smaller than that of *N_PI_* and was reversed. The interaction between *N_HD_* and the *Planting method* was not significant. DDSR and TPR represent the dry direct seeded rice and transplanted rice, respectively. AK and YA represent the cultivar’s names, ‘Akitakomachi’, and ‘Yumiazusa’, respectively. In our analysis of covariance, the regression lines were different between TPR and DDSR at the panicle initiation stage (left side, *P* = 0.0002) and heading stage (right side, *P* < 0.0001).

The difference in the spikelet-N relationship between planting methods and between growth stages could be related to the different crop N uptake patterns. We, therefore, applied a multiple regression analysis to examine the relative contribution of N uptake on spikelet density at different growth stages (Table 4). We found that both N_PI_ and N_PI-HD_ significantly contributed to spikelet density, while the *Planting method* (DDSR or TPR), *Year* (2018, 2019, or 2020), or *Cultivar* (‘Akitakomachi’ or ‘Yumiazusa’) had no significant effects (Table 4, Model 1). A simplified model without these categorical variables (Model 2) demonstrated that the coefficient for N_PI_ was more than twice as large as that of N_PI-HD_. The differences in crop N uptake pattern and sensitivity of spikelet density to N uptake at different growth stages accounted for much of the variation in spikelet density due to the N application method, *Planting method*, *Year*, and *Cultivar*.

**Table 4.** The effects of nitrogen uptake at different growth stages (PI and HD), planting methods (DDSR and TPR), cultivars (AK and YA), and years (2018, 2019, and 2020) on spikelet density.

| Explanatory variables | Model 1 |  | Model 2 |  |
| --- | --- | --- | --- | --- |
|  | Coefficient | P-value | Coefficient | P-value |
| Intercept | 7639 | <.0001 | 8005 | <.0001 |
| $N_{PI}$ | 3268 | <.0001 | 3038 | <.0001 |
| $N_{PI-HD}$ | 1698 | <.0001 | 1753 | <.0001 |
| $N_{HD}$ | - | - | - | - |
| *Planting method* |  |  |  |  |
| (DDSR) | -51 | 0.9225 | - | - |
| (TPR) | 51 | - | - | - |
| *Cultivar* (AK) | -713 | 0.0686 | - | - |
| (YA) | 713 | - | - | - |
| *Year* (2018) | -1070 | 0.1311 | - | - |
| (2019) | -16 | - | - | - |
| (2020) | 1086 | - | - | - |
| Fitting criteria |  |  |  |  |
| Adjusted R<sup>2</sup> | 0.60 |  | 0.58 |  |
| RMSE | 3665 |  | 3746 |  |
| AICc | 1743 |  | 1742 |  |
| BIC | 1761 |  | 1751 |  |

Model 1 includes categorical variables: *planting methods*, *cultivars*, and *years* and Model 2 excludes them.
DDSR, dry direct-seeded rice; TPR, transplanted rice.
AK and YA mean the cultivar's names, 'Akitakomachi', and 'Yumiazusa', respectively.

### Crop N dynamics and model parameters

Crop N uptake increased almost exponentially until 100 DAS and linearly until maturity, with noted differences among environments (Figure 4 and S5). When crop N was plotted against AETI (Equations 5 and 6), the differences narrowed (Figure S5), thus confirming that crop N uptake patterns for the vegetative growth period can be well represented by the model parameters (*N_LA5_*, and *RNR*).

**Figure 4.**
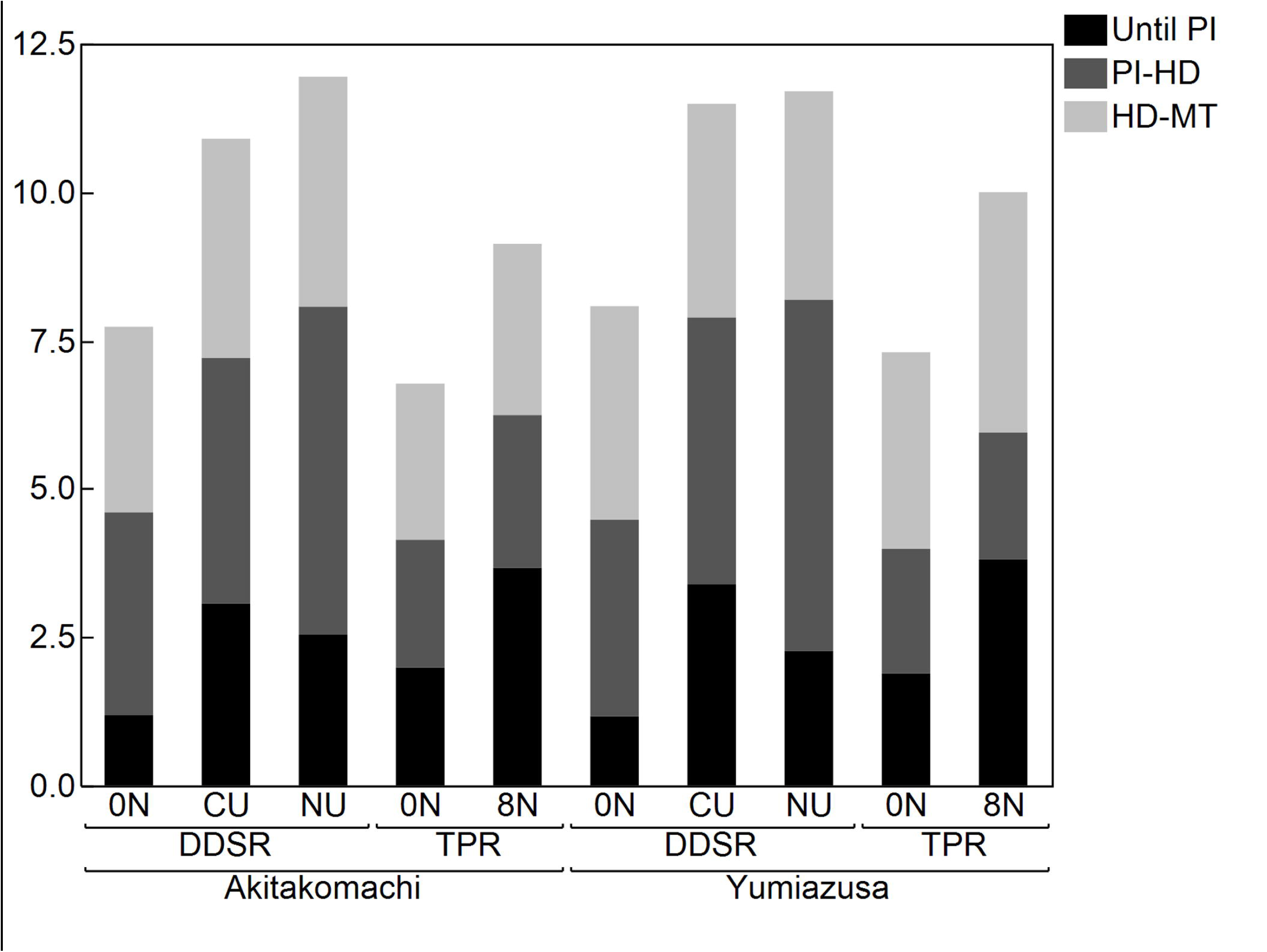
The relation between rice N uptake and DAS or AETI from LA5. Graph for YA-CU (Yumiazusa and Coated urea) treatment. Supplemental material (Figure S5) shows the relation of all treatments. DAS, days after sowing; AETI, accumulated effective thermal index (see material and method section). *N_up_* is the N uptake of the rice plant. NLA5 is the N uptake at 5th leaf age. RNR is the relative N uptake rate. NLA5 and RNR were derived as the intercept and coefficient of the exponent.

*N_LA5_* tended to be higher in CU, and that of 0N and NU had variation by year (Table 5). *RNR* had a different Y*N effect (*P*=0.0052) and tend to be higher in CU. YA had higher *N_LA5_* than AK while exhibiting similar *RNR* (Table 5).

**Table 5.**
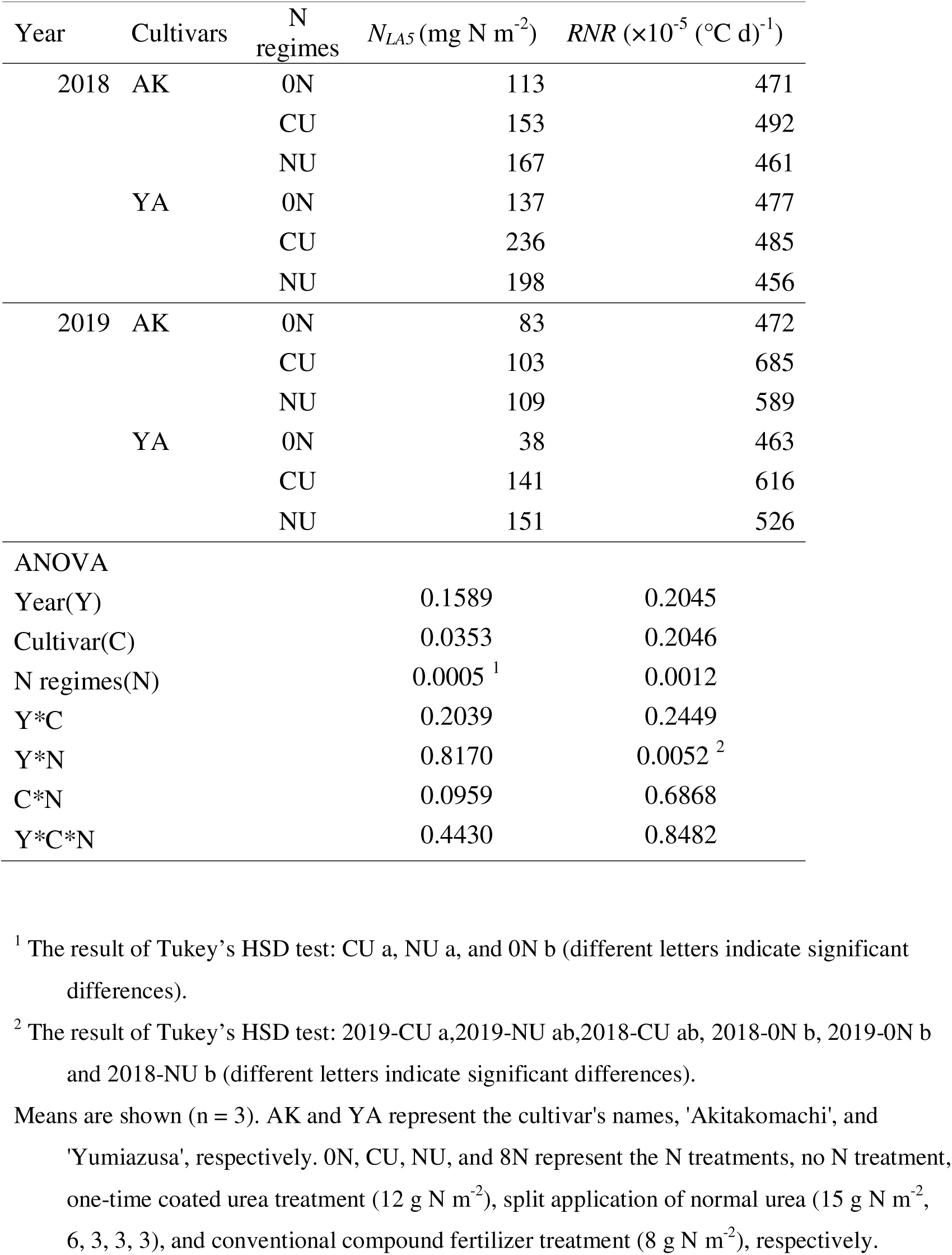
Nitrogen uptake at fifth leaf age (NLA5) and relative N uptake rate (RNR) of DDSR under different environments, nitrogen regimes, and cultivars.

In Model 3, adjusted R^2^ (0.85) was higher, besides AICc and BIC (66 and 72) being lower than in Model 4 which is the model that excludes *Year* (0.66, 95, and 100, respectively) (Table 6). The *Year* had a significant effect (*P =* 0.0099). The standardized partial regression coefficients (β) were 1.92 (*N_LA5_*) and 1.23 (*RNR*) in the model excluding the *Year* effect (Table 6). VIFs in Models 3 and 4 were >4, thus serious multicollinearity was not suggested between *N_LA5_* and *RNR* (Table 6).

**Table 6.**
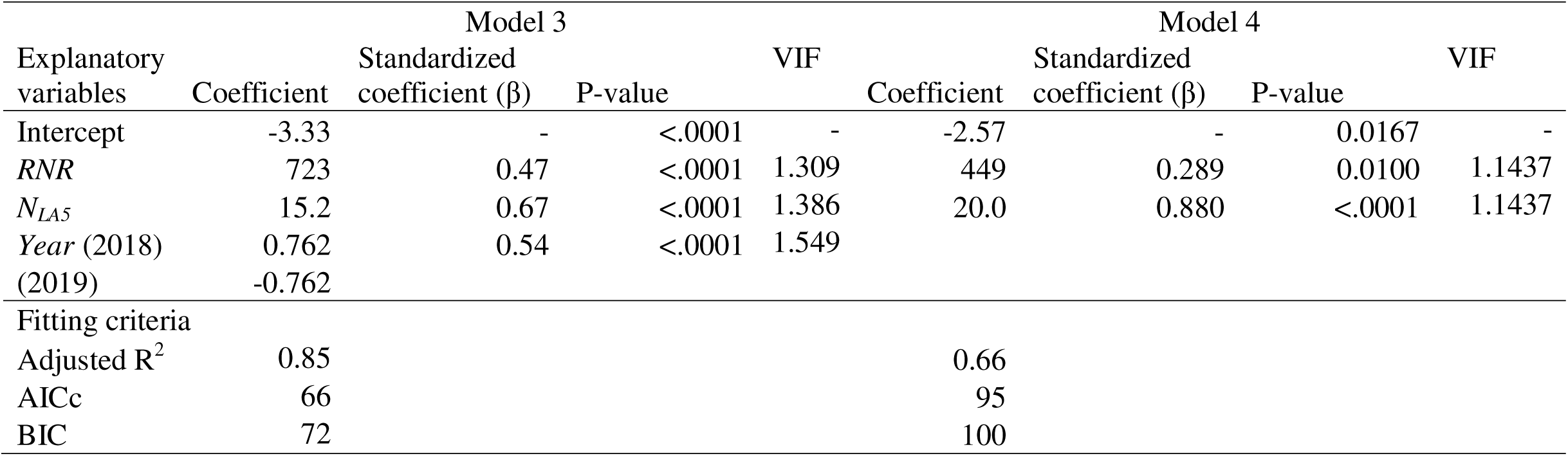
Multiple regression of N_PI_ using RNR, N_LA5_ and Year (2018 or 2019) as nitrogen uptake parameters (Model 3), and without year terms (Model 4) in DDSR.

## Discussion

Our three-year field experiment confirmed that DDSR yield was lower than TPR by an average of 11% (Table 2), similar to that reported by Shinoto et al., (2021).

Furthermore, the ANUE was lower in the DDSR than in the TPR by 33% (Table 3), which is in line with the observations that DDSR requires a greater amount of fertilizer to be productive (Otani et al., 2013). In this section, we first explore possible reasons for the low ANUE in DDSR compared to TPR. Then we discuss how N regimes and cultivars affect production efficiency in DDSR cultivation.

### Why is the ANUE of DDSR low? (Comparison with TPR)

ANUE can be considered a product of fertilizer N recovery and the efficiency of converting crop N into grain yield. We predicted that the former, represented by ANR, was largely responsible for the lower ANUE in DDSR relative to TPR because a large fertilizer loss was expected for the DDSR, particularly in an Andosol with a high percolation rate (Kanmuri et al., 2017). However, our analysis indicated that both ANR and CNUE were lower in the DDSR than those in the TPR, both contributing to the low ANUE of DDSR.

The low CNUE of DDSR was mainly induced by two factors. One was spikelet production per unit of *N_HD_* or *N_MT_* was limited in the DDSR relative to the TPR. As shown in Figure 2, *N_HD_* or *N_MT_* were comparable between DDSR and TPR, but the N uptake pattern was considerably different. While 8N of TPR took up 39% of the total crop N before the PI stage, CU of DDSR only 29% at PI while accelerating the N uptake toward heading and ultimately surpassing the N uptake observed in the TPR at the heading stage (Table 3 and Figure 2). The multiple regression analysis of spikelet density on *N_PI_* and *N_PI-HD_* revealed that the contribution of *N_PI_* was more than twice as large as that of *N_PI-HD_* (Table 4), suggesting that DDSR having a smaller *N_PI_* and larger *N_PI-HD_* leads to low spikelet production efficiency per unit N uptake. These results indicate that enhancing *N_PI_* might be effective in increasing CNUE and ANUE. Alternatively, Model 1 showed that the *N_PI_* and *N_PI-HD_* relationship was not significantly influenced by the planting method (DDSR or TPR), thus allowing us to show the universality of the relationship between spikelet density and N uptake for rice grown using different planting methods.

Secondly, the unfavorable climatic conditions for DDSR during the grain filling period compared with TPR; thus, limiting source capacity for grain growth. DDSR heading occurred approximately 9–17 days later than that of TPR, resulting in lower solar radiation and air temperature during the grain-filling period for DDSR plants. This is reflected in the poor grain filling (%) and 1000-grain weight at all N levels (Figure 1). Therefore, optimum growing seasons for DDSR need to be determined, accounting for cultivar phenological properties and site-specific climatic conditions.

### How can we improve N_PI_? (Comparison among treatments on DDSR)

N uptake before the PI stage is critical for yield determination and that is more important for DDSR than that for TPR as shown above section. We analyzed the N uptake pattern on DDSR until PI using the two parameters (*N_LA5_* and *RNR*) with the comparison of CU, NU, and 0N. The parameters of the N uptake pattern significantly influenced the *N_PI_* with *N_LA5_* and *RNR* exhibiting higher impacts (Table 6), thus suggesting that N uptake during the early stage, even N uptake at LA5, affects the yield.

The N application before LA5 plays an important role in leaf development and contributes to biomass and yield production. The days of LA5 were consistently earlier in order to CU, NU, and 0N (Table 1), suggesting the N application effect for initial leaf development. These initial differences in leaf development accounted for more than half of the differences in the final leaf number (Figure S2). The delay in leaf development may decrease the opportunity of booting and causes lower yield.

The ANR observed in this study was generally lower for DDSR than TPR (Table 3), which highlights the need for improvements regarding N application methods for DDSR. The ANUE associated with the NU treatment was significantly lower than that associated with the CU treatment (Table 3). This indicates that the lower yield of the NU treatment might have been caused by the imperfect application regimes rather than differences in recovery. Lower N_PI_ in NU than in CU also suggested that imperfection of the NU regime during the vegetative growth period, which includes the initial growth stage as noted above paragraph.

YA (high-yielding cultivar) retained its yield advantage under dry direct-seeding practice. The YA brown rice yield was significantly higher than that of the AK (Table 2). Model 1 for spikelet determination showed weak differences between AK and YA (Table 4, *P =* 0.0686), indicating YA may exhibit slightly higher CNUE than AK. YA had comparatively more panicles and 1000-grain weight but significantly lower grain-filling than AK; thus, causing a lower YA yield than AK yield in 2019 (Table S5). The DDSR in a cool climate may not provide YA with the appropriate conditions to express its high-yielding abilities because the grain-filling is restricted by unfavorable air temperature or solar radiation, especially at the grain-filling stage. Further investigations are needed for DDSR yield constraints associated with a meteorological condition in a cool climate.

### Remaining issues on DDSR in a cool climate

We have clarified two important matters to increase ANUE on DDSR in a cool climate: lower N_PI_ than that of TPR and the importance of initial N uptake (*N_LA5_*). However, It remains some questions: the possibility of increasing *N_LA5_*, and, as already noted above, meteorological constraints at the grain-filling stage in a cool climate. In NU treatment, it seems to be room for an increase N_LA5_ (Table 1). But normal urea fertilizer at an early stage may be lost for the most part by ammonia volatilization (Griggs et al., 2007; Qi et al., 2012) and denitrification(Casey et al., 2018) on DDSR. Further study is needed to increase *N_LA5_* by NU to the same level as that of CU.

Although some challenges remain to determine optimal N proportion and application stage in the split application of NU, the yield of NU treatment was not a significant difference compared to conventional CU treatment in this study. Therefore, NU treatment in this study indicated the possibility to be an alternative method to coated urea. Additionally, NU treatment could be a cheaper N fertilization method by their cheaper cost than that of CU though the labor cost for the topdressing need to examine.

## Conclusions

Our three-year field experiment confirmed that DDSR yield was lower than TPR by an average of 11%. ANR and CNUE were lower in the DDSR than in the TPR, contributing to the low ANUE of DDSR. The low CNUE of DDSR was mainly induced by lower N_PI_ and spikelet production per unit of *N_HD_* or *N_MT_* was limited in the DDSR relative to the TPR. We analyzed the N uptake pattern on DDSR until PI using the two parameters (*N_LA5_* and *RNR*) with the comparison of CU and NU. The parameters of the N uptake pattern significantly influenced the *N_PI_* with *N_LA5_* and *RNR* exhibiting higher impacts (Table 6), thus suggesting that N uptake during the early stage, even N uptake at LA5, affects the yield. The days of LA5 were consistently earlier in order to CU, NU, and 0N (Table 1), suggesting the N application effect for initial leaf development. Although some challenges remain to determine optimal N regimes in the split application of NU, NU treatment in this study indicated the possibility to be an alternative method to CU when it cannot be used.

## Disclosure statement

No potential conflict of interest was reported by the authors.

## Acknowledgements

We thank Hisaya Tanaka, Hiroshi Kawamukai, Eisaku Kumagai, Fumihiko Saito, and Kafumi Segawa for their help with in-field management and sampling. Professor Hiroyuki Shimono of Iwate University provided valuable suggestions. The colleagues of NARO Tohoku Agricultural Research Center and the United Graduate School of Agricultural Sciences-Iwate University supported our study.

## Funding

This work was supported by the commissioned project (high-technology development project; “Daikibo suiden einou”) of the Ministry of Agriculture, Forestry and Fisheries, Japan.

## Notes

### Competing Interest Statement

The authors have declared no competing interest.

### Summary of Updates

Analysis method, results, discussion and abstract were revised.

