## Supplementary material for "Nitrogen uptake pattern of dry direct-seeding rice and its contribution to yield in northeastern Japan": Figure S5

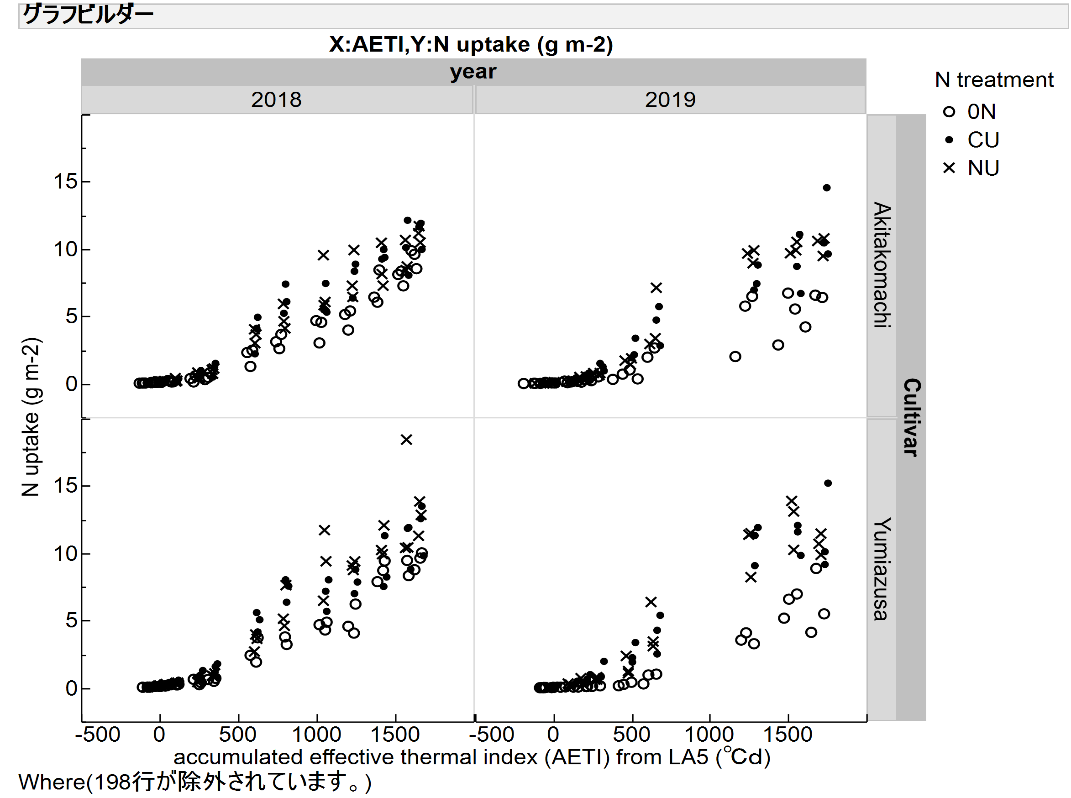

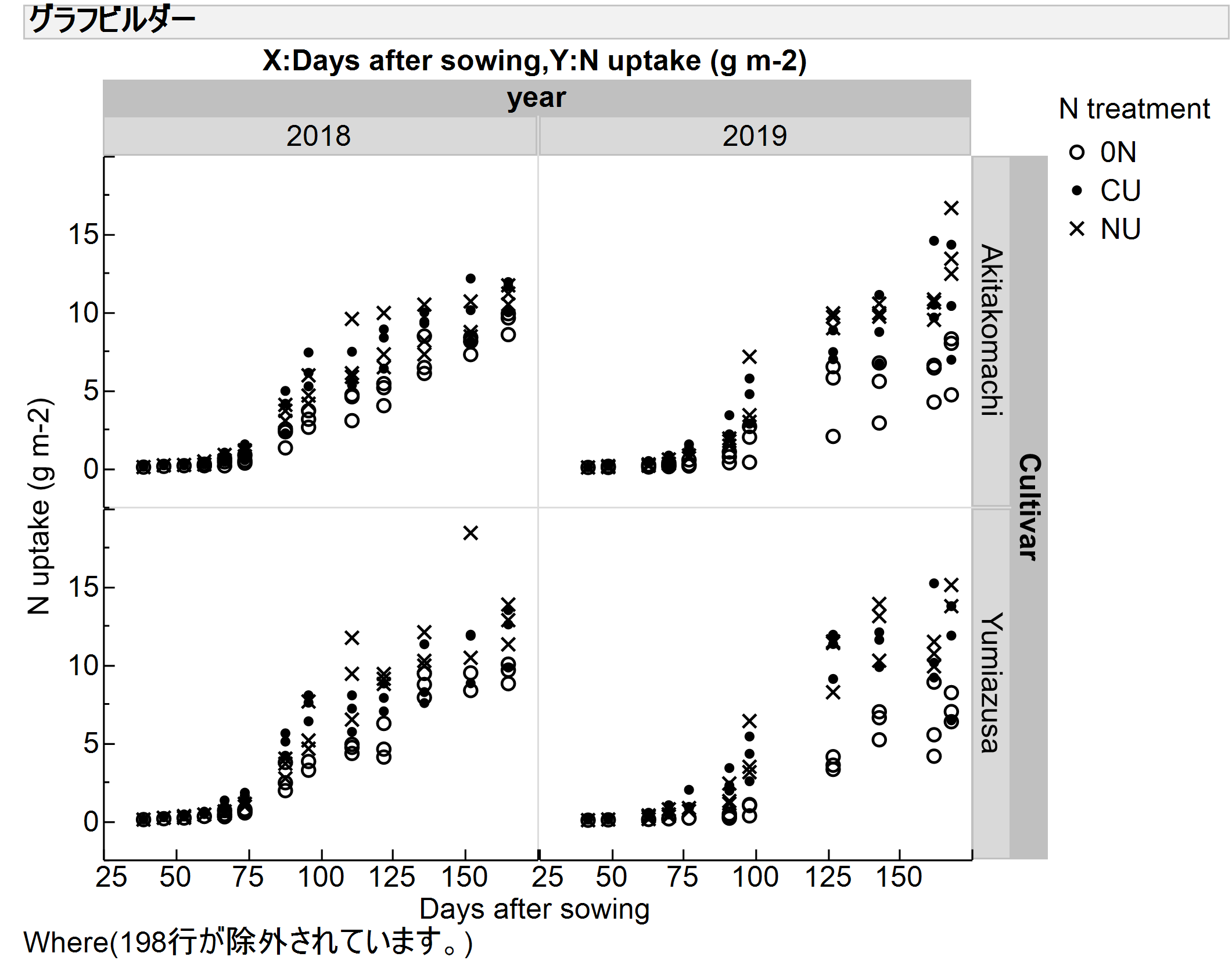

Figure S5. The relation between rice N uptake and days after sowing or accumulated effective thermal index (AETI) from LA5 in DDSR.

These graphs represent treatments conducted in this study.

0N, CU, and NU represent the N treatments, no N treatment, coated urea treatment, and split application of normal urea treatment, respectively.
