## Supplementary material for "Nitrogen uptake pattern of dry direct-seeding rice and its contribution to yield in northeastern Japan": Table S2

Table S2. The effective air temperature coefficient (Hanyu and Uchijima, 1962) for calculating AETI (accumulated effective thermal index) (Takahashi et al., 1976). The daily effective thermal index is a daily average of the product of the hourly air temperature and effective air temperature coefficient

| The class of temperature | Air temperature range(°C)^1^ | Effective air temperature coefficient |
| --- | --- | --- |
|  | ~10 | 0 |
| 11 | 10.1~12.0 | 0.045 |
| 13 | 12.1~14.0 | 0.170 |
| 15 | 14.1~16.0 | 0.300 |
| 17 | 16.1~18.0 | 0.420 |
| 19 | 18.1~20.0 | 0.523 |
| 21 | 20.1~22.0 | 0.619 |
| 23 | 22.1~24.0 | 0.715 |
| 25 | 24.1~26.0 | 0.809 |
| 27 | 26.1~28.0 | 0.900 |
| 29 | 28.1~30.0 | 0.976 |
| 31 | 30.1~32.0 | 1.000 |
| 33 | 32.1~34.0 | 1.000 |

1 Air temperature range was obtained by rounding to two decimal places.
