## Supplementary material for "Nitrogen uptake pattern of dry direct-seeding rice and its contribution to yield in northeastern Japan": Table S3

Table S3. The crop calendar of experiments (CU for DDSR and 8N for TPR)

| Year | Planting method | Cultivars | Emergence | | LA2^2^ | PI^3^ | HD^4^ | Harvest^5^ | |
| --- | --- | --- | --- | --- | --- | --- | --- | --- | --- |
|  |  |  | Date | DAS^1^ | DAS | DAS | DAS | Date | DAS |
| 2018 | DDSR | AK | 15-May | 18 | 27 | 96 | 116 | 12-Oct | 168 |
|  |  | YA | 15-May | 18 | 27 | 91 | 117 | 22-Oct | 178 |
|  | TPR | AK | - | - | - | 72 | 104 | 21-Sep | 149 |
|  |  | YA | - | - | - | 72 | 106 | 25-Sep | 153 |
| 2019 | DDSR | AK | 13-May | 21 | 24 | 91 | 114 | 7-Oct | 168 |
|  |  | YA | 13-May | 21 | 24 | 92 | 116 | 12-Oct | 173 |
|  | TPR | AK | - | - | - | 77 | 105 | 23-Sep | 156 |
|  |  | YA | - | - | - | 77 | 107 | 27-Sep | 160 |
| 2020 | DDSR | AK | 12-May | 20 | 21 | 92 | 120 | 6-Oct | 182 |
|  |  | YA | 12-May | 20 | 21 | 92 | 119 | 13-Oct | 182 |
|  | TPR | AK | - | - | - | 74 | 103 | 29-Sep | 159 |
|  |  | YA | - | - | - | 74 | 107 | 1-Oct | 161 |

^1^ Day after sowing.

^2^ LA2 is the date when leaf age 2 (perspective survey).

^3^ Panicle initiation date is defined as the date when the average panicle length of the main stem reached 2 mm.

^4^ Heading date is defined as the date when 50% heading of all panicles in the CU treatment.

^5^ Harvest was done after the grain yellowing rate reached 85%.
