## Supplementary material for "Nitrogen uptake pattern of dry direct-seeding rice and its contribution to yield in northeastern Japan": Table S4

Table S4. Brown rice yield and yield components for dry-direct seeded rice under different environments, nitrogen regimes, and cultivars

| Year | Cultivar | N  regimes | Brown rice  yield^1^  (g m^-2^) | | Panicle  density  (m^-2^) | | Spikelet  per panicle  (m^-2^) | | Spikelet  density  (10^3^ m^-2^) | | Grain  Filling (%) | | 1000-grain  weight (g) | |
| --- | --- | --- | --- | --- | --- | --- | --- | --- | --- | --- | --- | --- | --- | --- |
| 2018 | AK | 0N | 299 | | 268 | | 65.7 | | 17.5 | | 82.3 | | 22.3 | |
|  |  | CU | 448 | | 347 | | 76.1 | | 26.3 | | 78.1 | | 21.5 | |
|  |  | NU | 370 | | 326 | | 66.6 | | 21.7 | | 82.5 | | 22.5 | |
|  | YA | 0N | 349 | | 312 | | 63.8 | | 19.8 | | 79.0 | | 21.7 | |
|  |  | CU | 460 | | 452 | | 62.6 | | 28.2 | | 73.5 | | 22.0 | |
|  |  | NU | 470 | | 399 | | 63.9 | | 25.5 | | 77.3 | | 22.8 | |
| 2019 | AK | 0N | 192 | | 218 | | 56.7 | | 12.3 | | 90.0 | | 21.1 | |
|  |  | CU | 384 | | 331 | | 68.5 | | 20.9 | | 91.3 | | 21.7 | |
|  |  | NU | 403 | | 354 | | 67.6 | | 23.9 | | 85.3 | | 21.9 | |
|  | YA | 0N | 173 | | 231 | | 52.8 | | 12.2 | | 78.8 | | 21.1 | |
|  |  | CU | 383 | | 323 | | 72.1 | | 22.3 | | 85.0 | | 21.9 | |
|  |  | NU | 356 | | 396 | | 65.2 | | 24.7 | | 77.8 | | 21.4 | |
| 2020 | AK | 0N | 289 | | 247 | | 74.0 | | 18.6 | | 92.7 | | 20.9 | |
|  |  | CU | 478 | | 338 | | 81.0 | | 27.1 | | 92.4 | | 21.1 | |
|  |  | NU | 495 | | 369 | | 86.0 | | 31.3 | | 91.0 | | 21.1 | |
|  | YA | 0N | 381 | | 242 | | 73.9 | | 17.8 | | 87.0 | | 22.2 | |
|  |  | CU | 559 | | 351 | | 87.8 | | 30.9 | | 90.7 | | 22.2 | |
|  |  | NU | 503 | | 316 | | 81.0 | | 25.4 | | 89.5 | | 22.5 | |
| ANOVA | Year (Y) | | | 0.0174 | | 0.4615 | | 0.0134 | | 0.1164 | | 0.0319 | | 0.2036 |
|  | Cultivar (C) | | | 0.0083 | | 0.0937 | | 0.4709 | | 0.1104 | | 0.0002 | | 0.0043 |
|  | N regimes (N) | | | <.0001 ^2^ | | <.0001 ^2^ | | 0.0223 ^3^ | | <.0001 ^2^ | | 0.6829 | | 0.0163 ^4^ |
|  | Y*C | | | 0.0062 | | 0.0473 | | 0.6277 | | 0.0797 | | 0.2060 | | 0.0003 |
|  | Y*N | | | 0.0465 | | 0.3189 | | 0.6687 | | 0.4659 | | 0.0695 | | 0.0880 |
|  | C*N | | | 0.7393 | | 0.8453 | | 0.9465 | | 0.3614 | | 0.6904 | | 0.6232 |
|  | Y*C*N | | | 0.1158 | | 0.6211 | | 0.6769 | | 0.4148 | | 0.8952 | | 0.3340 |

^1^ Grain thicker than 1.85 mm was counted to calculate brown rice yield. Brown rice yield and 1000-grain weight are presented at a water content of 15%.

^2^ The result of Tukey’s HSD test: CU a, NU a, and 0N b (different letters indicate significant differences).

^3^ The result of Tukey’s HSD test: CU a, NU ab, and 0N b (different letters indicate significant differences)

^4^ The result of Tukey’s HSD test: NU a, CU ab, and 0N b (different letters indicate significant differences)

AK and YA represent the cultivar's names, 'Akitakomachi' and 'Yumiazusa'.

0N, CU, and NU represent the N treatments, no N treatment, coated urea treatment (12 g N m^-2^), and split application of normal urea (15 g N m^-2^) respectively.
