## Supplementary material for "Nitrogen uptake pattern of dry direct-seeding rice and its contribution to yield in northeastern Japan": Table S5

Table S5. Brown rice yield and yield components for transplanted rice under different environments, nitrogen regimes, and cultivars

| Year | Cultivar | N regimes | Brown rice yield^1^ | Panicle density | Spikelet per panicle | Spikelet density | Grain Filling | 1000-grain weight | Reference |
| --- | --- | --- | --- | --- | --- | --- | --- | --- | --- |
|  |  |  | (g m^-2^) | (m^-2^) | (m^-2^) | (10^3^ m^-2^) | (%) | (g) |  |
| 2018 | AK | 0N | 311 | 268 | 60.7 | 16.2 | 93.7 | 23.0 | Yabiku et al, 2021 |
|  |  | 8N | 481 | 360 | 67.5 | 24.3 | 93.7 | 22.7 |  |
|  | YA | 0N | 356 | 257 | 71.7 | 18.4 | 93.8 | 23.2 |  |
|  |  | 8N | 550 | 359 | 75.3 | 27.1 | 93.4 | 22.9 |  |
| 2019 | AK | 0N | 322 | 244 | 65.9 | 16.1 | 95.9 | 22.5 |  |
|  |  | 8N | 539 | 382 | 65.7 | 25.1 | 95.2 | 22.2 |  |
|  | YA | 0N | 383 | 269 | 71.7 | 19.3 | 95.0 | 22.9 |  |
|  |  | 8N | 602 | 391 | 78.1 | 30.5 | 95.8 | 22.9 |  |
| 2020 | AK | 0N | 312 | 242 | 61.0 | 14.7 | 92.9 | 22.2 |  |
|  |  | 8N | 431 | 308 | 65.8 | 20.2 | 93.8 | 22.1 |  |
|  | YA | 0N | 360 | 246 | 68.2 | 16.7 | 93.7 | 22.4 |  |
|  |  | 8N | 457 | 274 | 74.6 | 20.5 | 94.8 | 22.5 |  |

^1^Grain thicker than 1.85 mm was counted to calculate brown rice yield. Brown rice yield and 1000-grain weight are presented at a water content of 15%.

AK and YA represent the cultivar's names, 'Akitakomachi' and 'Yumiazusa'.

0N and 8N represent the N treatments, no N treatment and conventional treatment (8 g N m^-2^), respectively.
