## Supplementary material for "Nitrogen uptake pattern of dry direct-seeding rice and its contribution to yield in northeastern Japan": Table S6

Table S6. Rice plant nitrogen uptake and apparent nitrogen recovery rate of dry-direct seeded rice under different environments, nitrogen regimes, and cultivars

| Year | Cultivar | N regimes | N uptake (g m^-2^) | | | ANR | ANUE  (g g-N^-1^) | CNUE  (g g-N^-1^) | CNUEF  (g g-N^-1^) |
| --- | --- | --- | --- | --- | --- | --- | --- | --- | --- |
|  |  |  | PI | HD | MT |  |  |  |  |
| 2018 | AK | 0N | 2.0 | 4.6 | 9.3 | - | - | 32 | - |
|  |  | CU | 3.7 | 7.1 | 11.2 | 0.15 | 12.1 | 40 | 78 |
|  |  | NU | 3.6 | 7.6 | 11.1 | 0.12 | 5.0 | 33 | 39 |
|  | YA | 0N | 2.7 | 4.9 | 9.5 | - | - | 37 | - |
|  |  | CU | 4.9 | 7.5 | 12.0 | 0.21 | 10.8 | 38 | 44 |
|  |  | NU | 3.4 | 9.0 | 12.7 | 0.21 | 8.9 | 37 | 38 |
| 2019 | AK | 0N | 0.7 | 4.0 | 6.9 | - | - | 28 | - |
|  |  | CU | 2.5 | 6.2 | 10.6 | 0.30 | 16.3 | 36 | 52 |
|  |  | NU | 1.7 | 7.1 | 11.8 | 0.48 | 14.4 | 34 | 43 |
|  | YA | 0N | 0.3 | 3.7 | 6.7 | - | - | 26 | - |
|  |  | CU | 2.5 | 8.3 | 11.0 | 0.34 | 17.9 | 35 | 49 |
|  |  | NU | 1.6 | 8.4 | 11.6 | 0.50 | 12.6 | 31 | 37 |
| 2020 | AK | 0N | 0.9 | 5.2 | 7.0 | - | - | 41 | - |
|  |  | CU | 3.0 | 8.4 | 10.9 | 0.42 | 16.3 | 44 | 48 |
|  |  | NU | 2.4 | 9.5 | 12.9 | 0.50 | 15.9 | 38 | 35 |
|  | YA | 0N | 0.5 | 4.9 | 8.0 | - | - | 48 | - |
|  |  | CU | 2.7 | 7.8 | 11.5 | 0.37 | 15.6 | 49 | 51 |
|  |  | NU | 1.8 | 7.1 | 10.8 | 0.24 | 9.4 | 47 | 44 |
|  | ANOVA | Year (Y) | 0.0044 | 0.1329 | 0.1972 | 0.0182 | 0.0674 | 0.1198 | 0.3416 |
|  |  | Cultivar (C) | 0.9612 | 0.5134 | 0.4943 | 0.9040 | 0.8113 | 0.0825 | 0.1827 |
|  |  | N regimes (N) | <.0001 ^1^ | <.0001 ^2^ | <.0001 ^2^ | 0.4603 | 0.0023 | 0.0009 ^3^ | 0.0268 |
|  |  | Y*C | 0.1058 | 0.0317 | 0.4484 | 0.1856 | 0.2711 | 0.0201 | 0.1864 |
|  |  | Y*N | 0.9884 | 0.8797 | 0.1475 | 0.3445 | 0.4527 | 0.1193 | 0.8111 |
|  |  | C*N | 0.4388 | 0.6126 | 0.5739 | 0.5126 | 0.5637 | 0.866 | 0.0942 |
|  |  | Y*C*N | 0.8022 | 0.2421 | 0.2316 | 0.6058 | 0.2934 | 0.8615 | 0.0371 |

^1^ The result of Tukey’s HSD test: CU a, NU b, and 0N c (different letters indicate significant differences).

^2^ The result of Tukey’s HSD test: CU a, NU a, and 0N b (different letters indicate significant differences).

^3^ The result of Tukey’s HSD test: CU a, NU b, and 0N b (different letters indicate significant differences)

DDSR, dry direct seeding cultivation; TPR, transplanted cultivation.

AK and YA represent the cultivar's names, 'Akitakomachi' and 'Yumiazusa', respectively.0N, CU, and NU represent the N treatments, no N treatment, coated urea treatment (12 g N m^-2^), and split application of normal urea (15 g N m^-2^) treatment, respectively.
