## Supplementary material for "Nitrogen uptake pattern of dry direct-seeding rice and its contribution to yield in northeastern Japan": Figure S2

Difference in leaf number at seeding stage1

| Difference in final leaf number from CU | 0N |
| --- | --- |
| -0.888889 | -0.23 |
| -0.444444 | -0.266667 |
| -1.66667 | -1.13333 |
| -1.11111 | -0.944444 |

| Difference in final leaf number from CU | NU |
| --- | --- |
| -0.111111 | -0.0666667 |
| -0.222222 | -0.333333 |
| 0 | -0.677778 |
| -0.222222 | -0.877778 |

Figure S2. The relationship between the leaf number differences in the CU treatment condition at the seedling stage and the final leaf number.

0N, CU and NU represent the N treatments, no N treatment, coated urea treatment, and split application of normal urea, respectively.

The regression line is based on 0N treatment plots.

^1^ The seedling stage was at 45 (2018), and 49 (2019) days after sowing, respectively.
