## Supplementary material for "Nitrogen uptake pattern of dry direct-seeding rice and its contribution to yield in northeastern Japan": Figure S3

### Crop development and biomass growth of DDSR

Tiller number started to increase at around LA7 (or around 70 DAS) in all years and differences among N treatments became apparent (Figure S3). The 0N plot had consistently fewer tillers than the CU and NU plots but the difference was smaller in 2018 than in 2019. Of the two fertilized plots, CU had more tillers than NU in 2018, but in 2019, NU produced more tillers than CU for both cultivars from around 80 DAS onward. The Tiller number in CU started to increase from LA6 (around 40 DAS), whereas the onset of tillering in NU was delayed until LA7 (50 DAS) when the second topdressing was applied. Subsequently, the tiller number increased sharply in the NU treatment and surpassed CU from 75 DAS onward.

Aboveground biomass started to increase at around 75 DAS and almost linearly toward maturity in 2018, whereas it peaked at around 135 DAS in 2019 (Figure S4). The 0N treatment had smaller biomass almost for the entire growing season than that of the other two treatments, while the difference between the CU and NU treatments was not as apparent as in tiller number (Figure S3). The final aboveground biomass in 2018 tended to be greater than the other year.

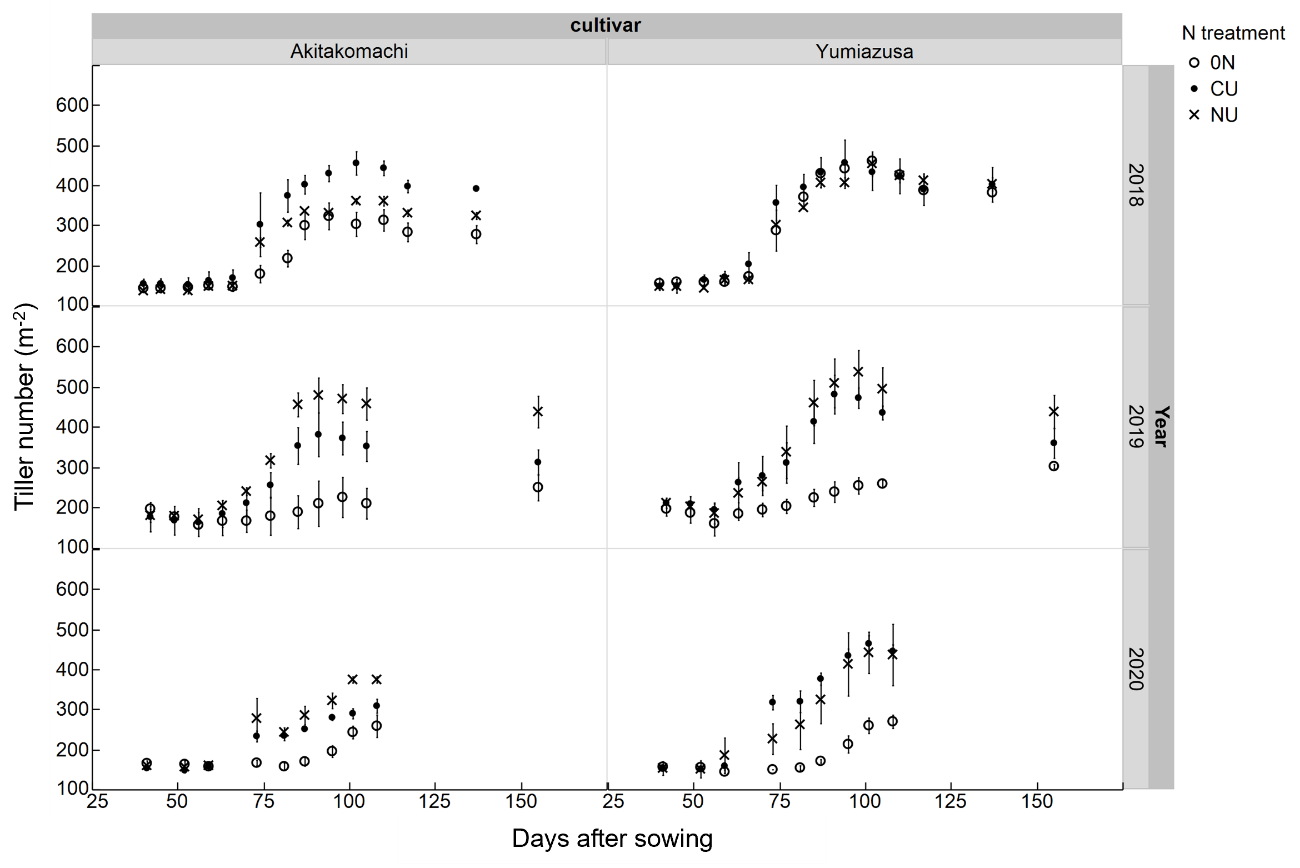

Figure S3. Tiller number of dry direct-seeded rice under different environments, nitrogen regimes, and cultivars.

The bars indicate a standard error of three replicate samples.

0N, CU, and NU represent the N treatments, no N treatment, coated urea treatment (12 g N m^-2^), and split application of normal urea (15 g N m^-2^), respectively.

For the NU treatment, a total of 15g N m^-2^ of urea was split-applied four times at 40, 66, 82, and 94 days after sowing (DAS) in 2018, 45, 66, 80, 94 DAS in 2019, and 52, 67, 80, 95 DAS in 2020, at a dose of 6 g N m^-2^, 3 g N m^-2^, 3 g N m^-2^, and 3 g N m^-2^, respectively.
