## Supplementary material for "Nitrogen uptake pattern of dry direct-seeding rice and its contribution to yield in northeastern Japan": Figure S4

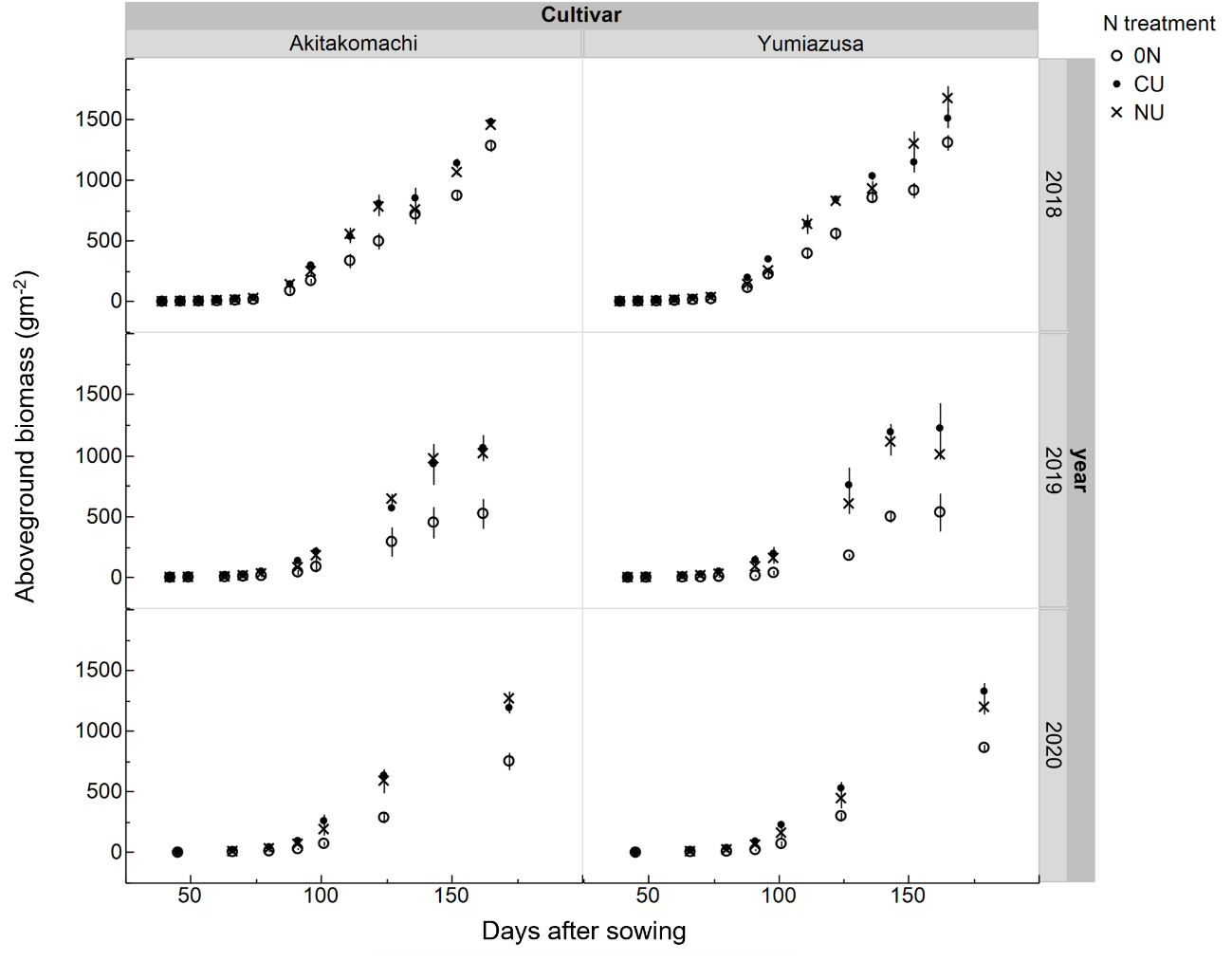

Figure S4. Aboveground biomass of dry direct-seeded rice under different environments, nitrogen regimes, and cultivars.

The bars indicate a standard error of three replicate samples.

0N, CU, and NU represent the N treatments, no N treatment, coated urea treatment (12 g N m-2), and split application of normal urea (15 g N m-2), respectively.

For the NU treatment, a total of 15g N m-2 of urea was split-applied four times at 40, 66, 82, and 94 days after sowing (DAS) in 2018, 45, 66, 80, 94 DAS in 2019, and 52, 67, 80, 95 DAS in 2020, at a dose of 6 g N m-2, 3 g N m-2, 3 g N m-2, and 3 g N m-2, respectively.
